# Molecularly defined subpopulations of leptin receptor neurons dissociate the control of food intake from blood pressure

**DOI:** 10.64898/2026.03.26.714551

**Authors:** Allison M. Duensing, Dylan Belmont-Rausch, Abigail J. Tomlinson, Aiden Crowley, Frederike Sass, Elizabeth Heaton, Bernd Coester, Jenny M. Brown, Shad Hassan, Zhe Wu, Nathan Qi, David P. Olson, Paul V. Sabatini, Martin G. Myers, Tune H. Pers

## Abstract

While previous studies have suggested that leptin regulates cardiovascular function independently of body weight, the specific leptin receptor (*Lepr)*-expressing neurons that mediate these distinct effects remain unknown. We found that genes located in blood pressure (BP)-associated genome-wide association study loci were regulated by leptin in *Lepr* and glucagon-like peptide-1 receptor (*Glp1r*)-expressing (Lepr^Glp1r^) neurons. Ablating *Lepr* from these cells decreased BP despite causing hyperphagic obesity. Single-cell and spatial transcriptomics revealed that Lepr^Glp1r^ neurons segregate into two distinct subpopulations of cells located in the arcuate nucleus (ARC) and dorsomedial hypothalamic nucleus (DMH). Activating ARC Lepr^Glp1r^ neurons suppressed food intake without impacting energy expenditure or cardiovascular function. Conversely, DMH Lepr^Glp1r^ neurons increased energy utilization and BP without altering food intake. Our results identify distinct Lepr^Glp1r^ neuron subpopulations that dissociate the control of food intake from outputs related to sympathetic tone, including BP, suggesting the potential therapeutic utility of targeting of these subpopulations independently.

## Introduction

The prevalence of obesity continues to drive the development of its comorbidities, including cardiovascular diseases [1]. The accumulation of fat mass increases circulating concentrations of leptin, a hormone secreted by adipose tissue in proportion to body energy stores. Leptin acts on specialized leptin receptor (LepRb, encoded by *Lepr*)-expressing neurons in the hypothalamus to decrease food intake and enable energy expenditure via increased sympathetic nervous system (SNS) output and neuroendocrine function [2, 3].

The lack of leptin (or melanocortin 4 receptor (*MC4R*), which contributes to leptin action) decreases blood pressure (BP) and cardiovascular risk despite promoting extreme obesity [4–6]. Furthermore, increased leptin concentrations correlate with increased BP and brown adipose tissue (BAT) thermogenesis [7, 8] independently of body weight. Hence, in addition to suppressing food intake and body weight, leptin increases BP and other SNS-associated variables independently of its effects on adiposity.

Consistent with the pleiotropic effects of leptin, the hypothalamus contains multiple distinct populations of *Lepr* neurons, each with presumably different functions [2, 9]. For instance, the arcuate nucleus (ARC) contains multiple populations of *Lepr* neurons that control food intake, including *Agrp* neurons promote food seeking and suppress energy expenditure [10, 11] and *Pomc* neurons that decrease feeding and augment energy utilization. *Lepr* neurons of the dorsomedial hypothalamic nucleus (DMH) control SNS outflow to BAT, the cardiovascular system, and other effectors [7, 8]. Identifying the mechanisms by which leptin controls BP and other cardiovascular parameters and separating these from the leptin-mediated control of food intake and energy balance has the potential to identify new therapeutic targets for cardiometabolic diseases.

Unfortunately, the *Lepr* neurons that modulate SNS outflow have not been defined molecularly, and direct leptin action on *Agrp* and *Pomc* neurons only modestly impacts food intake and body weight. In contrast, *Glp1r*-expressing *Lepr* neurons (Lepr^Glp1r^ neurons, which also express *Trh* [12] and *Bnc2* [13]) reside in a continuum across the ARC and DMH and play important roles in the control of feeding and body weight via their inhibition of *Agrp* neurons [14–16].

To identify *Lepr* neuron populations mediating the control of BP by leptin, we correlated the expression levels of genes associated with BP in genome-wide association studies (GWAS) [17] and leptin-regulated transcriptional programs in individual populations of hypothalamic *Lepr* neurons. This analysis implicated Lepr^Glp1r^ neurons in BP control. Consistently, ablating *Lepr* from these cells decreased BP in addition to promoting hyperphagic obesity. Furthermore, we found that Lepr^Glp1r^ neurons consist of two subpopulations of neurons with distinct transcriptional and anatomic features. These populations also perform different functions-one controls food intake while the other controls BP and other processes modulated by SNS outflow. Hence, these results not only define the molecular identity of a neuron population that mediates the control of BP by leptin but also segregate this BP circuits from the control of food intake.

## Results

### Enrichment of human BP-associated genes in the transcriptional response to leptin in BP-controlling Lepr^Glp1r^ neurons

To define cell type-specific transcriptional responses to leptin in individual populations of hypothalamic neurons, we used data from a parallel study [18]where we examined the transcriptional response to leptin treatment across mediobasal hypothalamic (MBH) cell populations from wild-type mice, including five *Lepr*-expressing cell types (Figure 1a, b).

**Figure 1.**
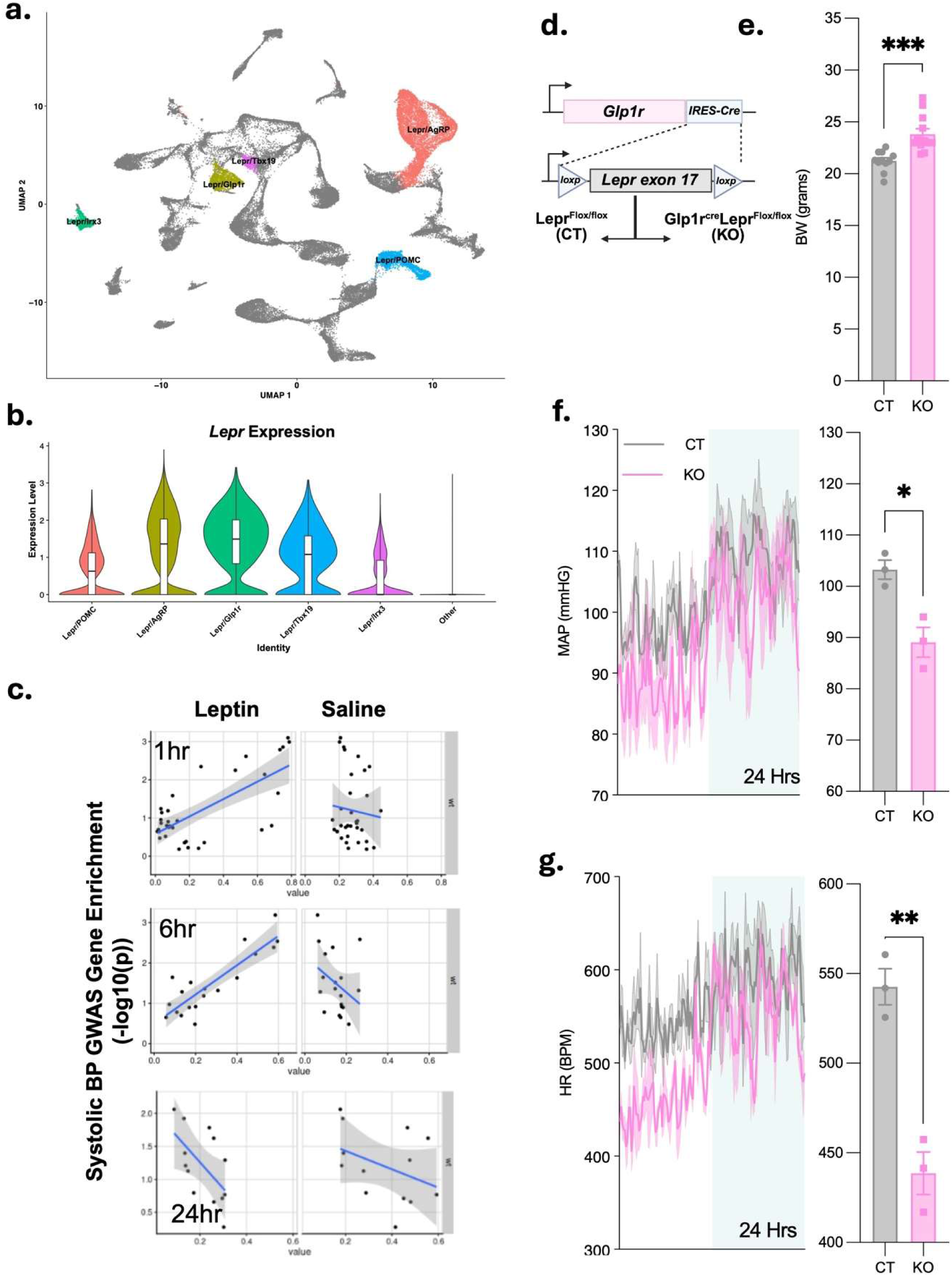
Roles for Lepr^Glp1r^ neurons in the control of cardiovascular function by leptin. (a) UMAP plot showing MBH neuronal cell types identified during snRNA-seq analysis of tissue from mice treated with saline or exogenous leptin [18]. *Lepr*-expressing population are denoted by labels and colors. (b) *Lepr* expression for the 5 identified MBH *Lepr* cell types, along with all other cell types. (c) Correlation of systolic BP-associated GWAS genes with the transcriptional response to leptin (left panels) or saline (right panels) for Lepr^Glp1r^ neurons from WT mice. (d) Schematic showing generation of *Lepr^Flox/Flox^* control (CT) and *Glp1r^Cre^;Lepr^Flox/Flox^* (Lepr^Glp1r^KO; KO) mice. (e) Body weight for CT and KO animals (n=11 CT and 12 KO). (f, g) Mean arterial pressure (MAP; f) and HR (g) over 24 hours (left panel) and averaged over the first 30 minutes of the light cycle (right panel) for CT (n=3) and KO (n=3) females. \**P* < 0.05 by Welche’s *t* test.

To identify which of these neuron populations might contribute to the control of BP by leptin, we correlated the expression of genes within systolic BP GWAS loci [17] to their likelihood of being regulated by leptin in each of the five MBH neuron cell populations (Figure 1c). We found a significant correlation between GWAS-defined genes and leptin-regulated genes only in Lepr^Glp1r^ neurons, after one (*P*=2.48x10^-6^), three (*P=*1.72x10^-3^) and six (*P=*1.37x10^-6^) hours of leptin treatment. This correlation was lost by 24 hours after treatment, however, consistent with the expected attenuation of exogenous leptin action.

Given the concordance between BP-associated genes and leptin responses in Lepr^Glp1r^ neurons, we next sought to understand the potential contribution of Lepr^Glp1r^ neurons to the control of BP. We therefore generated *Glp1r^Cre^;Lepr^Flox/Flox^* (Lepr^Glp1r^KO) mice lacking *Lepr* specifically in Lepr^Glp1r^ neurons (Figure 1d). Lepr^Glp1r^KO mice weighed significantly more than control animals, consistent with their previously demonstrated hyperphagic obesity (Figure 1e) [19]. To measure BP in these animals, we implanted intra-aortic probes to enable the real-time telemetric monitoring of heart rate (HR) and BP. This analysis revealed Lepr^Glp1r^KO mice have decreased mean arterial pressure and HR at the onset of the light cycle, despite their obesity (Figure 1f, g). Thus, direct leptin action on Lepr^Glp1r^ neurons increases BP and HR, as predicted by our human genetics-informed computational analysis.

### Transcriptional and anatomical subpopulations of hypothalamic Lepr^Glp1r^ neurons

To better classify the transcriptional identity of Lepr^Glp1r^ neurons (as well as other hypothalamic *Lepr* neuron populations), we analyzed the transcriptomes of *Lepr* neurons (Figure 2; Supplemental Figures 1-2). To this end, we isolated hypothalamic tissue from *Lepr^Cre^;Rosa26^LSLSun1-sfGFP^* mice (n=4) (Lepr^Sun1-sfGFP^ mice, in which the nuclear envelope of *Lepr*-expressing cells contains a green fluorescent protein (GFP) variant) and collected GFP-positive nuclei. We subjected these nuclei to snRNA-seq, yielding 25,546 nuclei across the four samples. Of these, 23,424 passed quality control (Supplemental Figure 1), of which 23,060 (98.4%) represented neurons (Supplemental Figure 2), consistent with the known restriction of LepRb expression to neurons in the hypothalamus and the expression of Cre recombinase from the LepRb-specific *Lepr* exon in *Lepr^Cre^* [20].

**Figure 2.**
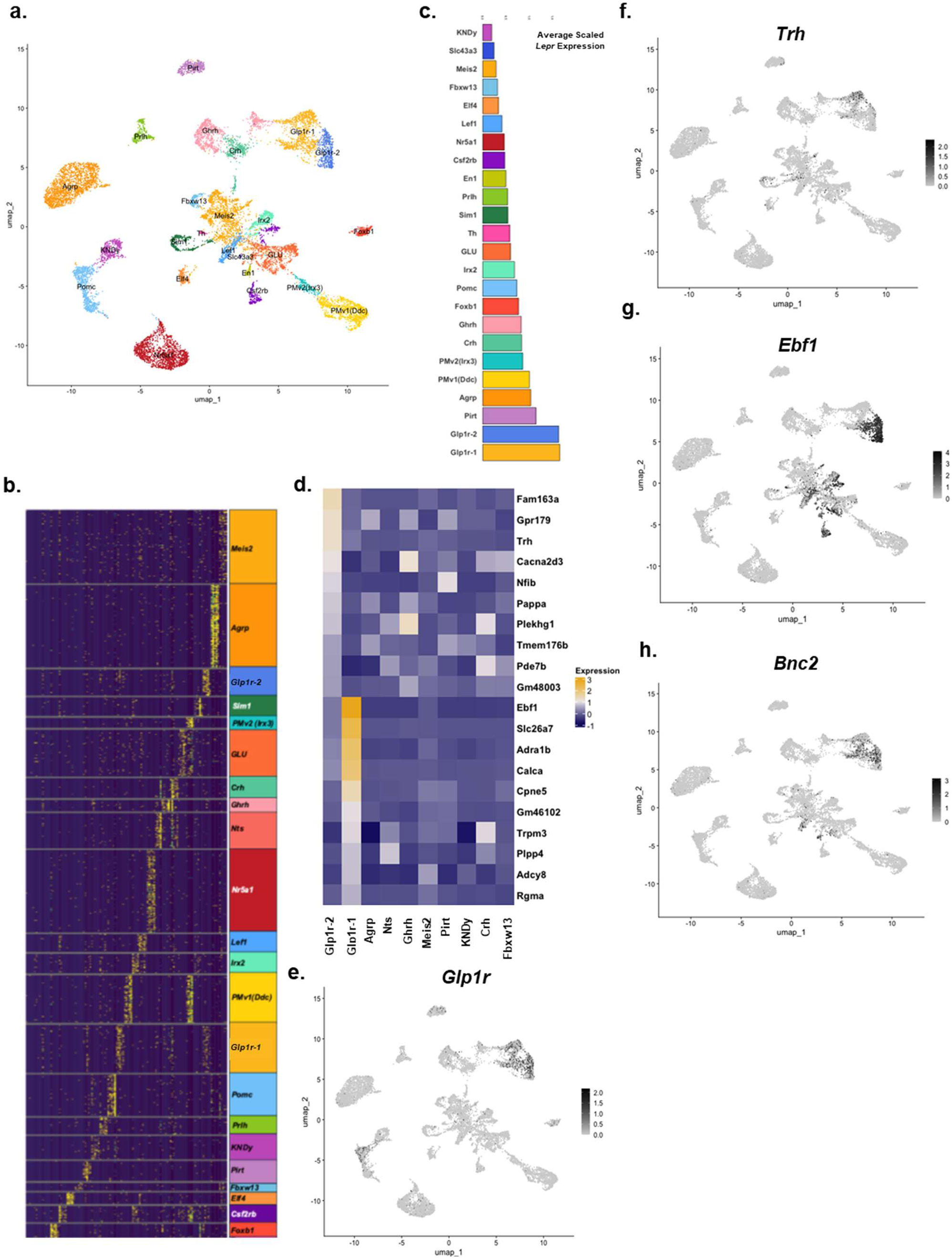
Hypothalamic *Lepr* neuron-specific snRNA-seq identifies novel populations of *Lepr* neurons, including distinct subpopulations of Lepr^Glp1r^ neurons. (a) UMAP projection of hypothalamic neuron clusters for GFP-expressing cells from Lepr^Sun1-sfGFP^ mice. (b) Heatmap of the top five genes per cluster with a log2 fold change > 2. (c) Average scaled *Lepr* expression for each neuron cluster. (d) Heatmap of gene expression the top 10 genes differentially expressed between Lepr^Glp1r-1^ and Lepr^Glp1r-2^ populations shown for all major GABAergic (*Slc32a1*-expressing) *Lepr* neuron clusters. Genes examined were all detected in >25% of the relevant cell cluster. (e-h) UMAP feature plots showing cells expressing *Glp1r* (e), *Trh* (f), *Ebf1* (g), *and Bnc2* (h); expression scales for the relevant gene shown to the right of each plot.

While our previous snRNA-seq analysis identified 18 MBH *Lepr* neuron populations [21], the increased resolution resulting from the relatively large number of nuclei in this analysis identified 24 distinct populations, including several that had not previously been detected (Figure 2a-b). Furthermore, some previously identified populations of *Lepr* neurons, including Lepr^Glp1r^ neurons, segregated into multiple distinct subpopulations in this new analysis. While both subpopulations of Lepr^Glp1r^ neurons (Lepr^Glp1r-1^ and Lepr^Glp1r-2^) neurons expressed *Glp1r*, *Slc32a1* (identifying them as GABAergic), and high levels of *Lepr*, many genes exhibited differential expression between the two Lepr^Glp1r^ cell types (Figure 2c-e; Supplemental Figure 2). Prominent differentially expressed genes between these two subpopulations included *Trh* (more highly expressed in Lepr^Glp1r-1^ neurons) and *Ebf1* (specific to Lepr^Glp1r-2^ neurons) (Figure 2d, f-g; supplemental figure 2). *Bnc2*, another marker for Lepr^Glp1r^ neurons, was distributed across both subpopulations (Figure 2h). Previous studies identified two *Trh*-expressing populations of ARC neurons, marked by the expression of *Cxcl12* or *Lef1*, respectively [12]. The *Trh*-expressing Lepr^Glp1r^ subpopulation coexpresses *Cxcl12*, not *Lef1* (Supplemental Figure 2).

We next generated a *Glp1r^Flp^* allele for use in combination with *Lepr^Cre^* (*Lepr^Cre^;Glp1r^Flp^* mice), thereby permitting the use of molecular genetic systems that require both Cre and Flp recombinases to manipulate Lepr^Glp1r^ neurons without impacting other anatomically interspersed classes of *Lepr* neurons. We initially bred *Lepr^Cre^;Glp1r^Flp^* mice onto a background containing alleles that mediate the Cre-dependent expression of GFP (*Rosa26^LSL-Gfp-L10a^*) and the Flp-dependent expression of tdTomato (*Rosa26^FSF-tdTomato^*), generating Lepr^GFP^;Glp1r^tdT^ reporter mice (Figure 3a). While this line underreported *Glp1r*-expressing cells (consistent with the decreased activity of Flp relative to Cre recombinase), the co-expression of the two fluorophores none-the-less permitted the identification of many Lepr^Glp1r^ neurons. These neurons lay in a continuum that spanned from the DMH to the ARC (Figure 3b-o), consistent with previous results [12, 13, 16, 19].

**Figure 3.**
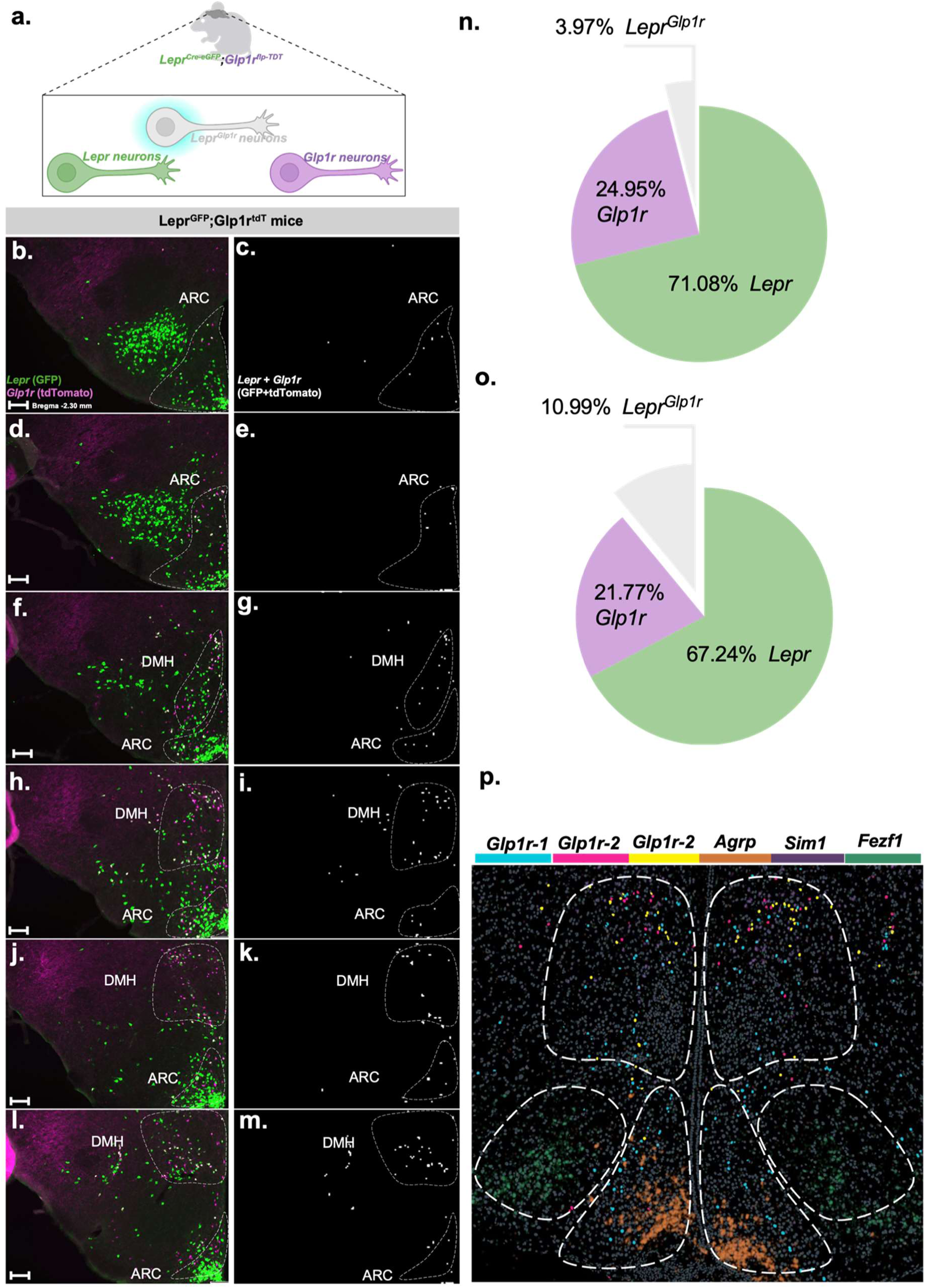
Distribution of Lepr^Glp1r^ neurons and their subtypes throughout the hypothalamus. (a) Schematic of Lepr^GFP^;Glp1r^tdT^ mouse model. (b-m) Representative images showing GFP (green) and tdTomato (red) (c, e, g, I, k, m) neurons or neurons containing colocalized GFP plus tdTomato (white) (d, f, h, j, l, n) throughout the Lepr^GFP^;Glp1r^tdT^ mouse hypothamalus (representative of n= 4 female and 6 male mice), running caudal (bregma -2.30; b,c) to rostral. Scale bars=100nm. (n) Mapping of Lepr^Glp1r-1^ (*Lepr+Glp1r+Trh*; blue) and Lepr^Glp1r-2^ (*Lepr+Glp1r+Ebf1*; red and *Lepr+Glp1r+Trh+Ebf1*; yellow) neurons onto a sagittal mouse section brain at the indicated rostral-caudal position in the hypothalamus. Distribution of *Sim1* (purple) and *Nr5a1* (green) signals is also shown.

To understand potential anatomic differences between the two subpopulations of Lepr^Glp1r^ neurons, we used spatial transcriptomics to define the distribution of each set of neurons (Figure 3p). This analysis revealed that while Lepr^Glp1r-1^ cells (which are *Ebf1*-negative) cluster in the ARC, Lepr^Glp1r-2^ cells (which contain *Ebf1*) reside primarily in the DMH. Hence, targeting Lepr^Glp1r^ cells in the ARC can enable the specific manipulation of the Lepr^Glp1r-1^ subpopulation, while targeting DMH Lepr^Glp1r^ cells can permit the specific manipulation of Lepr^Glp1r-2^ neurons.

We employed the Lepr^GFP^;Glp1r^tdT^ reporter mice in FOS studies to understand the regulation of each subpopulation of Lepr^Glp1r^ neurons *in vivo* (Supplemental Figure 4). Following an overnight fast, Lepr^Glp1r^ cells in both areas contained little detectable FOS (approximately 5% in the ARC and 6% in the DMH). Treatment with leptin (1 mg/kg, IP) or the GLP1R agonist, semaglutide (10 nmol/kg, IP) for 2 hours increased the number of FOS-containing Lepr^Glp1r^ cells in the ARC and DMH to approximately 15% (leptin) or 20% (semaglutide) – although the response to leptin was not statistically significant for the DMH cells. Hence, both ARC and DMH Lepr^Glp1r^ neurons exhibit low activity in food-deprived animals but respond to leptin and GLP1R agonists, as expected from their expression of the receptors for these ligands.

### Divergent Circuitry for ARC and DMH Lepr^Glp1r^ cells

To define the direct neural inputs to each subpopulation of Lepr^Glp1r^ cells, we developed a RAbies Dual Recombinase Retrograde (RADRR) tracing system (Figure 4a). This system utilizes the co-injection of twin helper viruses-one AAV that mediates the Cre-dependent expression of TVA (the cell surface receptor for the EnvA-pseudotyped rabies virus) and another AAV to mediate the Flp-dependent expression of the rabies glycoprotein (G; required for rabies packaging and retrograde viral propagation). The previously described EnvA-psuedotyped Rabies^ΔG-GFP^ virus [22] is subsequently injected at the same site, several weeks after helper AAV injection.

**Figure 4.**
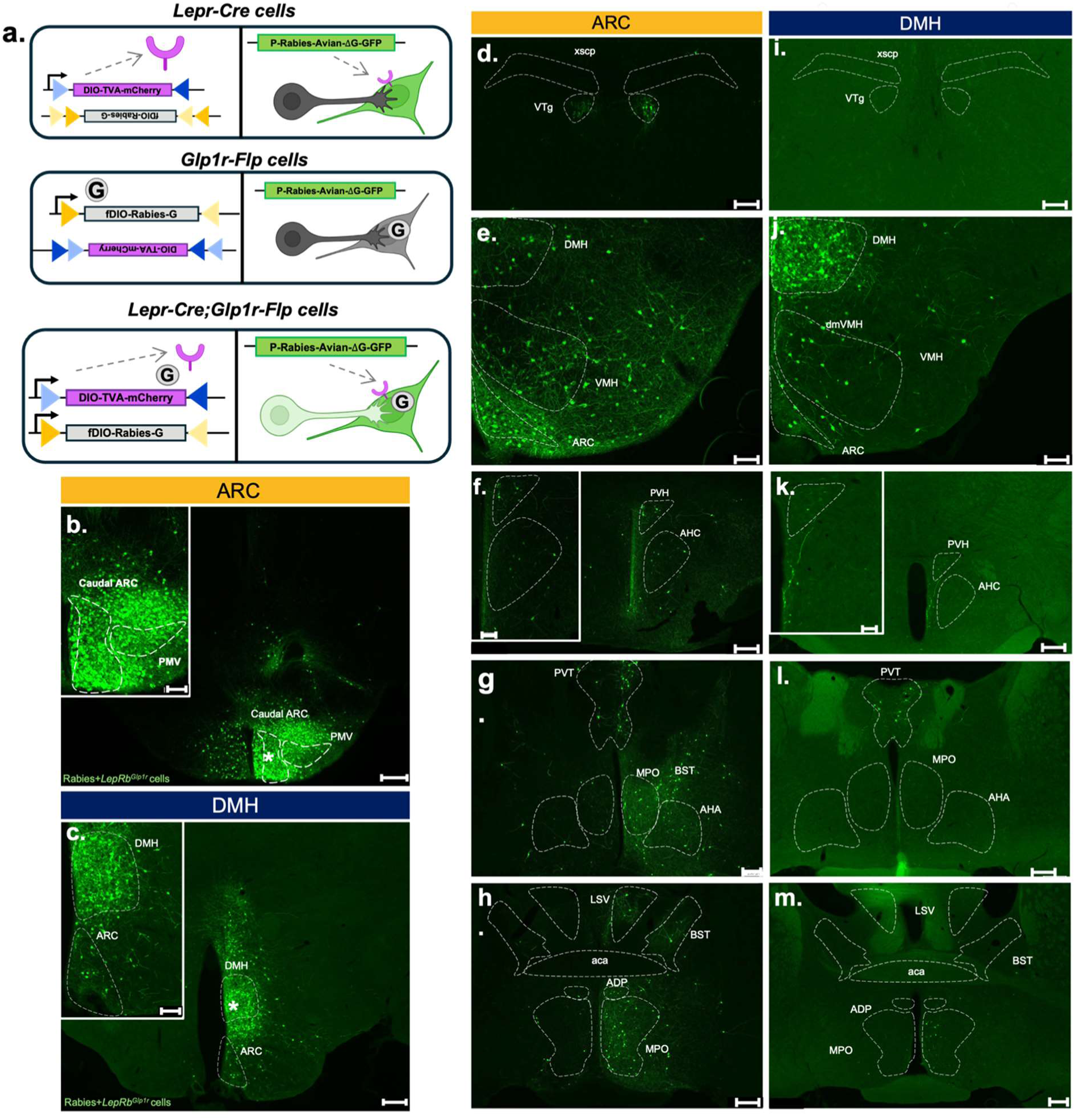
<u>RA</u>bies <u>D</u>ual <u>R</u>ecominase <u>R</u>etrograde (RADRR) system-mediated tracing from ARC and DMH Lepr^Glp1r^ neurons. (a) Schematic of RADRR tracing system. Adeno-associated viruses (AAVs) containing rabies glycoprotein (G) AAV8-hSyn1-DIO-TVA-mCherry (purple plasmid) expresses receptor for EnvA (TVA) in *Lepr^Cre^* cells; AAV8-hSyn-fDIO-Rabies-G (grey plasmid) expresses rabies-G in *Glp1r^Flp^* cells. Pseudotyped Rabies-ΔG-GFP (green plasmid) infects all *Lepr^Cre^* cells and can propagate one synapse retrograde from *Lepr^Cre^;Glp1r^Flp^* cells. (b, c) Representative images showing Rabies-ΔG-GFP-infected cells (green) at the viral hit site (*) in the ARC (b, representative of n=1 female and n=3 males) and DMH (c, representative of n=2 females and n=2 males) in *Lepr^Cre^;Glp1r^Flp^* mice. (d-m) Representative images showing Rabies-ΔG-GFP-infected cells (green) in regions afferent to ARC (d-h) or DMH (i-m) Lepr^Glp1r^ cells. Shown are ventral tegmental area (VTA) (d, i), DMH (e, j), PVH, anterior hypothalamic nucleus (AHA) (f, k), periventricular thalamus (PVT), medial preoptic (MPO), and bed nucleus of the stria terminalis (BST) (g-m). Main images, scale bars= 200 um. Insets show digital zooms of relevant areas in main images; scale bars= 100um.

When employing this system in *Lepr^Cre^;Glp1r^Flp^* mice, pseudotyped Rabies^ΔG-GFP^ virus specifically enters TVA-expressing *Lepr^Cre^* cells, but can only spread retrogradely from the subset of these cells that also express Flp recombinase/Rabies G. Hence, in *Lepr^Cre^;Glp1r^Flp^* mice the pseudotyped Rabies^ΔG-GFP^ virus can only propagate retrogradely from Lepr^Glp1r^ cells. Note that because pseudotyped Rabies^ΔG-GFP^ virus can enter *Lepr* cells independently of whether they contain *Glp1r*, it is not feasible to examine local retrograde tracing with this system, however.

To confirm the dual recombinase specificity of RADRR, we delivered it into the DMH of *Lepr^Cre^* mice-which produced GFP expression exclusively within the injection site. Similarly, delivery of RADRR into *Glp1r^Flp^* mice produced no labelling (Supplemental Figure 4). In contrast, injecting RADRR into the ARC or DMH of *Lepr^Cre^;Glp1r^Flp^* mice resulted in the detection of many GFP-expressing (retrogradely labelled) cells outside of the injection site (Figure 4d-m).

Delivery of the RADRR tracing system to the ARC of *Lepr^Cre^;Glp1r^Flp^* mice identified upstream inputs to ARC Lepr^Glp1r^ neurons in the nearby VMH and PVH, as well as in the VTg, PVT, and in the MPO, BST, and other rostral regions (Figure 4d-h). Targeting RADRR to DMH Lepr^Glp1r^ cells revealed GFP-containing inputs in some regions that overlapped with those from ARC Lepr^Glp1r^ cells-including the VMH, PVH, and PVT (although VMH inputs to DMH Lepr^Glp1r^ neurons tended to lie more medially than the VMH inputs to ARC Lepr^Glp1r^ neurons) (Figure 4i-m). Additionally, DMH Lepr^Glp1r^ neurons received no input from the VTg and little input from neurons in the MPO, BST, or other rostral regions.

Note that while we detected afferents to DMH Lepr^Glp1r^ cells in the ARC and afferents to ARC Lepr^Glp1r^ neurons in the DMH, the inability of the RADRR system to distinguish local inputs from *Lepr^Cre^* neurons within the injection plume prevented us from determining whether DMH Lepr^Glp1r^ neurons receive inputs from the DMH and whether ARC Lepr^Glp1r^ neurons receive inputs from the ARC. These results indicated that although both ARC and DMH Lepr^Glp1r^ neurons are activated by leptin and semaglutide and both subpopulations receive input from some overlapping areas, each Lepr^Glp1r^ subpopulation also receives many distinct inputs, consistent with potentially different functions for ARC and DMH Lepr^Glp1r^ neurons.

To define the regions to which ARC and DMH Lepr^Glp1r^ neurons project, we injected the tTARGIT AAV system [23] into the ARC or DMH of *Lepr^Cre^;Glp1r^Flp^* mice to express ChR2-YFP specifically in Lepr^Glp1r^ neurons in these regions (Figure 5). Consistent with previous results and with the dual recombinase requirement for this system, we detected no ChR2-YFP expression following the injection of these viruses into mice that contained only *Lepr^Cre^* or only *Glp1r^Flp^* (Supplemental Figure 5).

**Figure 5.**
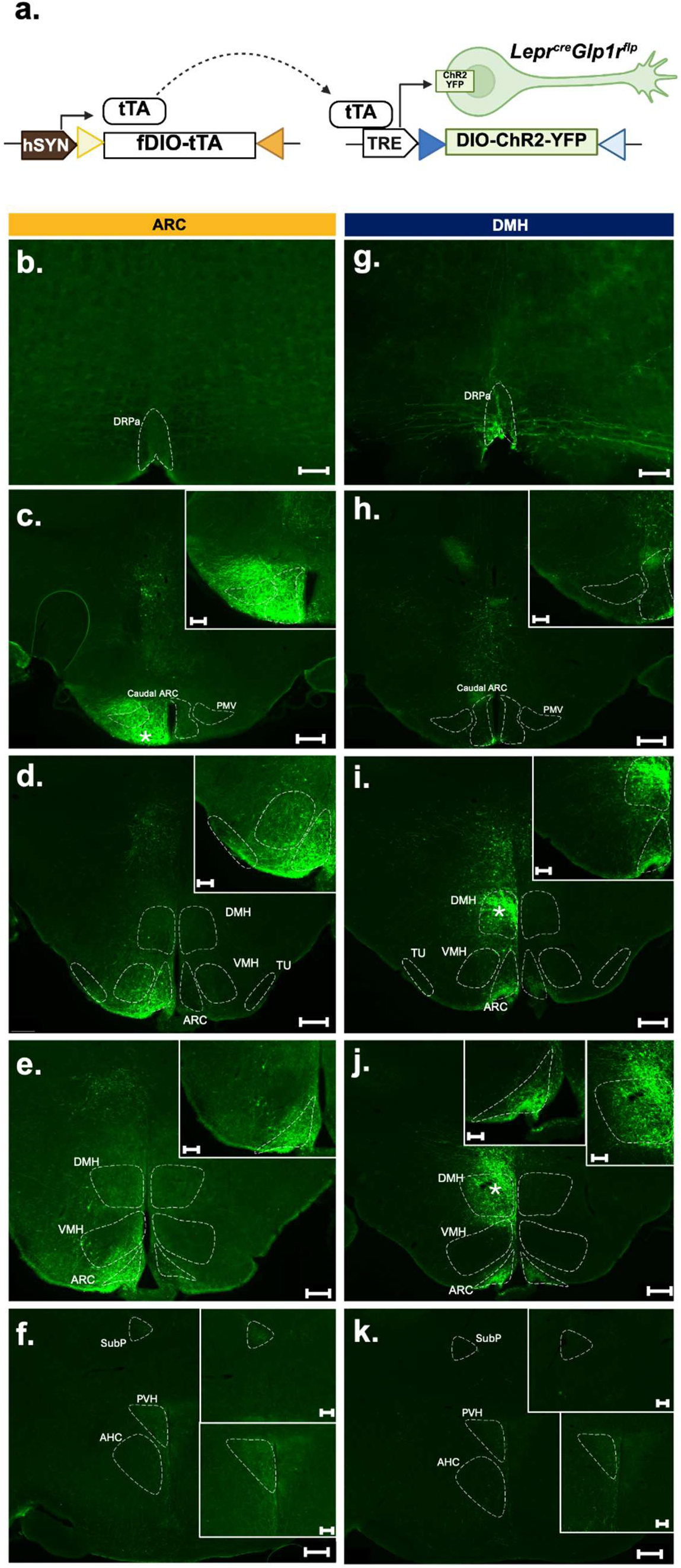
Downstream projections from ARC and DMH Lepr^Glp1r^ neurons. (a) Schematic of the tTA-driven Recombinase-Guided Intersectional Targeting (tTARGIT) viral system consisting of AAV8-SYN1-TETOX2-FLEX-FRT-tTA (grey plasmid) used to express tetracycline transactivator (tTA) in *Glp1r^Flp^* cells; tTA drives the expression of channelrhodopsin-2 (ChR2-eYFP) from rAAV8-TRE-DIO-hChR2-eYFP *Lepr^Cre^Glp1r^Flp^* cells. (b-k) Representative images showing GFP-immunoreactivity (green) in GFP-IR containing regions from *Lepr^Cre^Glp1r^Flp^* mice injected in the ARC (b-f, representative of n=2 males and 2 females) and DMH (g-k, representative of n=3 males). Viral hit site indicated by *. Scale bars in main images= 200 um. Scale bars in digital zoom inlay images= 100um.

Examining the YFP-containing projections from ChR2-YFP-expressing ARC and DMH Lepr^Glp1r^ neurons demonstrated strong projections into the mediobasal ARC (the location of *Agrp* neurons) for both cell types (Figure 5), consistent with their previously-described innervation (and inhibition) of *Agrp* neurons [14–16]. In contrast, only DMH Lepr^Glp1r^ neurons projected to the raphe pallidus (RPa; which modulates SNS outflow to cardiac and adipose tissues (among others)). Additionally, only ARC Lepr^Glp1r^ neurons sent dense projections to the tuberal nucleus (TU) and the PVH. Thus, while ARC and DMH Lepr^Glp1r^ neurons both project to the *Agrp* neuron-containing mediobasal ARC, each also targets unique brain regions.

### Divergent functions for ARC and DMH Lepr^Glp1r^ neurons

Although ARC and DMH Lepr^Glp1r^ neurons share many properties, the differences in their gene expression profiles, anatomic distribution, inputs, and projections suggest some divergence of function for these two cell types. To define potential differences in function for ARC and DMH Lepr^Glp1r^ subpopulations, we injected tTARGIT AAVs [8] into the ARC or DMH of *Lepr^Cre^;Glp1r^Flp^* mice to generate animals expressing activating (hM3Dq) designer receptors exclusively activated by designer drugs (DREADDs) [24] in ARC or DMH Lepr^Glp1r^ neurons (Lepr^Glp1r-ARC-Dq^ and Lepr^Glp1r-DMH-Dq^ mice, respectively), enabling their activation by the injection of the DREADD ligand, clozapine-N-oxide (CNO) (Figure 6a). As expected, injection of the Cre-dependent AAV-TRE-DIO-hM3Dq-mCherry cargo vector alone into *Lepr^Cre^;Glp1r^Flp^* mice in the absence of the Flp-dependent AAV-hSynfDIO-tTA driver virus failed to mediate substantial hM3Dq expression (Supplemental Figure 6), consistent with the dual recombinase specificity of this system.

**Figure 6.**
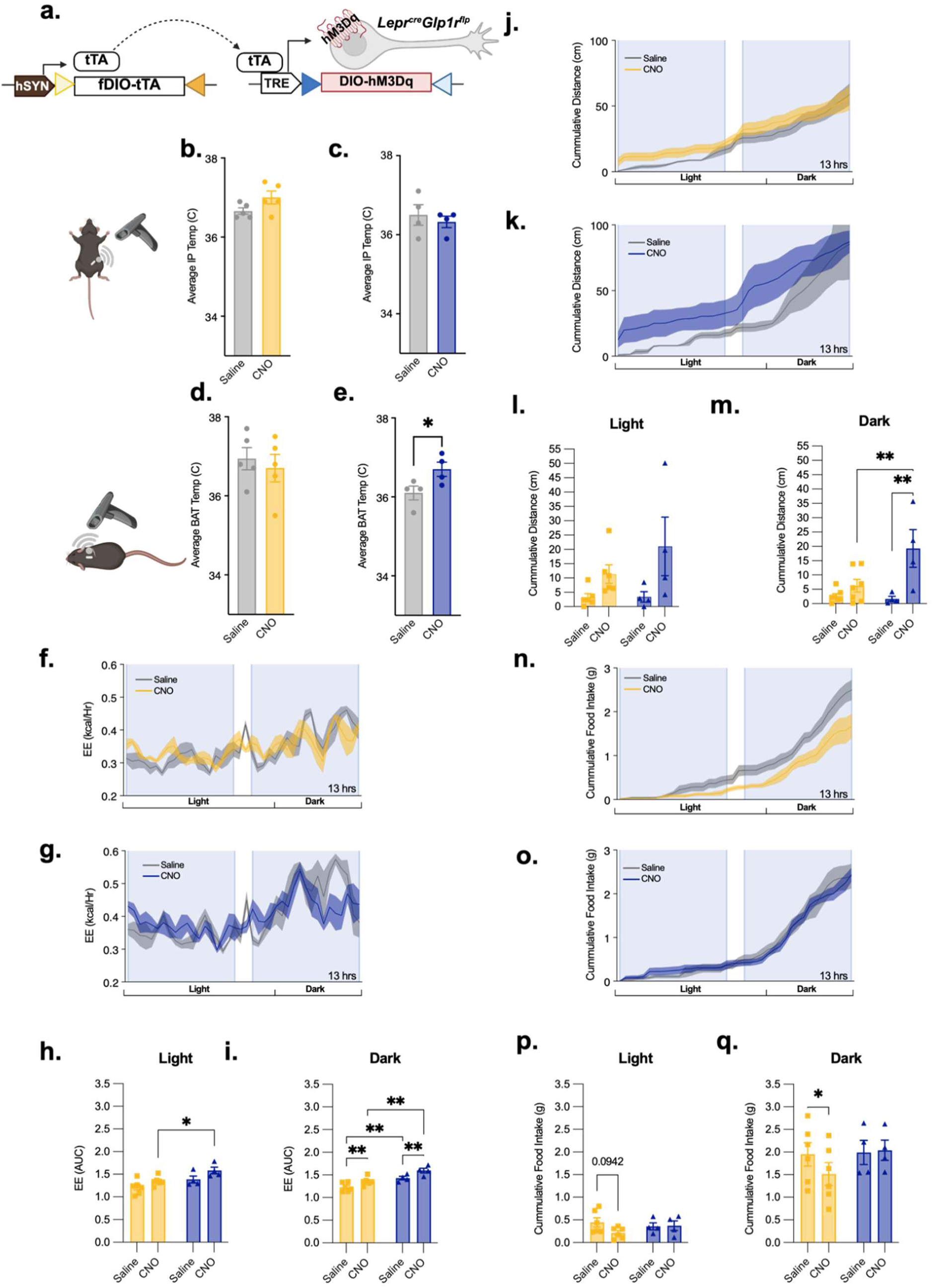
Modulation of food intake and energy expenditure by ARC and DMH Lepr^Glp1r^ neurons in female mice. (a) Schematic of the tTARGIT viral system used to express hM3Dq in Lepr^Glp1r^ neurons. AAV8-SYN1-TETOX2-FLEX-FRT-tTA (grey plasmid) mediates the expression of tTA in *Glp1r^Flp^* cells, which drives the expression of hM3Dq-mCherry from AAV8-TRE-DIO-hM3Dq-mCherry in *Lepr^Cre^Glp1r^Flp^* cells. (b, c) IP (core body) temperature (b, c) and intrascapular (BAT) temperature (d, e) over the 360 minutes following the activation of ARC (b, d, n=5) or DMH (c, e, n=4) Lepr^Glp1r^ neurons. Shown are averages +/-SEM; \**P* < 0.05 by Paired *t* test. (f-p) Continuous measurement of energy expenditure (f-i), cumulative locomotor activity (j-m), and cumulative food intake (n-q) following the injection of saline or CNO in Lepr^Glp1r-ARC-Dq^ (yellow; n=5) or Lepr^Glp1r-DMH-Dq^ mice (blue; n=4). Measurements were taken automatically using the SABLE metabolic cage system with 12 hour light and dark cycle, during which the animals were acclimated for 2 days, and then treated BID (0900 and 1600) with saline for 3 days and then with CNO (1 mg/kg, IP) for 3 days. Comparisons were made between the third day of saline injections and first day of CNO injections. (f, g, j, k, n, o) show continuous or cumulative measures of the indicated parameters over 13 hours; injections were performed at the beginning of the shaded areas. Graphs show summed values over the first hour (energy expenditure, locomotor activity) or six hours (food intake). Bar graphs shown averages +/-SEM; *p < 0.05, **p< 0.01 for the indicated comparisons by ANOVA with Fisher’s LSD post hoc test.

While we previously showed that Lepr^Glp1r^ neurons mediate the restraint of food intake by leptin, the finding that DMH Lepr^Glp1r^ neurons project to the RaP suggested that these cells might also contribute to the control of SNS outflow. Thus, we initially assessed the ability of activating ARC or DMH Lepr^Glp1r^ neurons to increase thermogenesis. We implanted temperature-monitoring telemetry chips into the peritoneal cavity or intrascapular region of Lepr^Glp1r-ARC-Dq^ and Lepr^Glp1r-DMH-Dq^ mice to monitor core body temperature or brown adipose tissue (BAT) thermogenesis, respectively (Figure 6b-e; Supplemental Figure 7). While core (IP) body temperature did not change in response to the activation of either ARC or DMH Lepr^Glp1r^ neurons (Figure 6b, c), activating DMH (but not ARC) Lepr^Glp1r^ neurons increased intrascapular temperature (Figure 6d, e), consistent with a specific role for DMH Lepr^Glp1r^ neurons in the control of BAT thermogenesis.

We next placed Lepr^Glp1r-ARC-Dq^ and Lepr^Glp1r-DMH-Dq^ animals in metabolic cages to measure additional SNS-related parameters, along with food intake, at baseline and during CNO administration (Figure 6f-q; Supplemental Figure 7). We found that activating DMH Lepr^Glp1r^ neurons near the onset of the light cycle or the dark cycle rapidly increased energy expenditure and locomotor activity in female and male mice, while ARC Lepr^Glp1r^ neurons produced no significant changes in these parameters (Figure 6f-m; Supplemental Figure 7). Thus, DMH Lepr^Glp1r^ neurons, but not ARC Lepr^Glp1r^ neurons, augment physiological and behavioral parameters associated with increased SNS outflow.

In contrast to the results above, we found that activating ARC Lepr^Glp1r^ neurons (but not DMH Lepr^Glp1r^ neurons) suppressed feeding during the dark cycle in female mice (these changes were not statistically significant in male mice, however) (Figure 6n-q; Supplemental Figure 7). Consistently, activating ARC Lepr^Glp1r^ neurons (but not DMH Lepr^Glp1r^ neurons) tended to decrease refeeding following an overnight fast (Supplemental Figure 8). Hence, activating ARC Lepr^Glp1r^ neurons suppresses food intake and activating DMH Lepr^Glp1r^ neurons augments SNS-associated parameters, but not vice-versa, consistent with distinct functions for each Lepr^Glp1r^ neuron subpopulation.

Our GWAS data analysis and *Lepr* knockout studies (Figures 1) predicted that Lepr^Glp1r^ neurons contribute to the control of BP and leptin action in the DMH contributes to the control of BP and other effects modulated by SNS outflow. Hence, we used telemetry in Lepr^Glp1r-ARC-Dq^ and Lepr^Glp1r-DMH-Dq^ mice to determine whether the control of cardiovascular function by Lepr^Glp1r^ neurons might also be specific to the DMH subpopulation (Figure 7). Indeed, we found that while activating ARC Lepr^Glp1r^ neurons tended to decrease (albeit not significantly) SNS-linked parameters of cardiovascular function (Figure 7a-d), activating DMH Lepr^Glp1r^ neurons increased systolic, diastolic, and mean arterial pressure as well as HR (Figure 7e-h). Hence, DMH Lepr^Glp1r^ neurons, but not ARC Lepr^Glp1r^ neurons, increase HR and BP. Overall, these findings demonstrate that the two subpopulations of Lepr^Glp1r^ neurons not only exhibit differences in gene expression, location, and circuitry, but also mediate distinct actions; DMH Lepr^Glp1r^ neurons augment SNS-linked effects, while ARC Lepr^Glp1r^ neurons suppress food intake.

**Figure 7.**
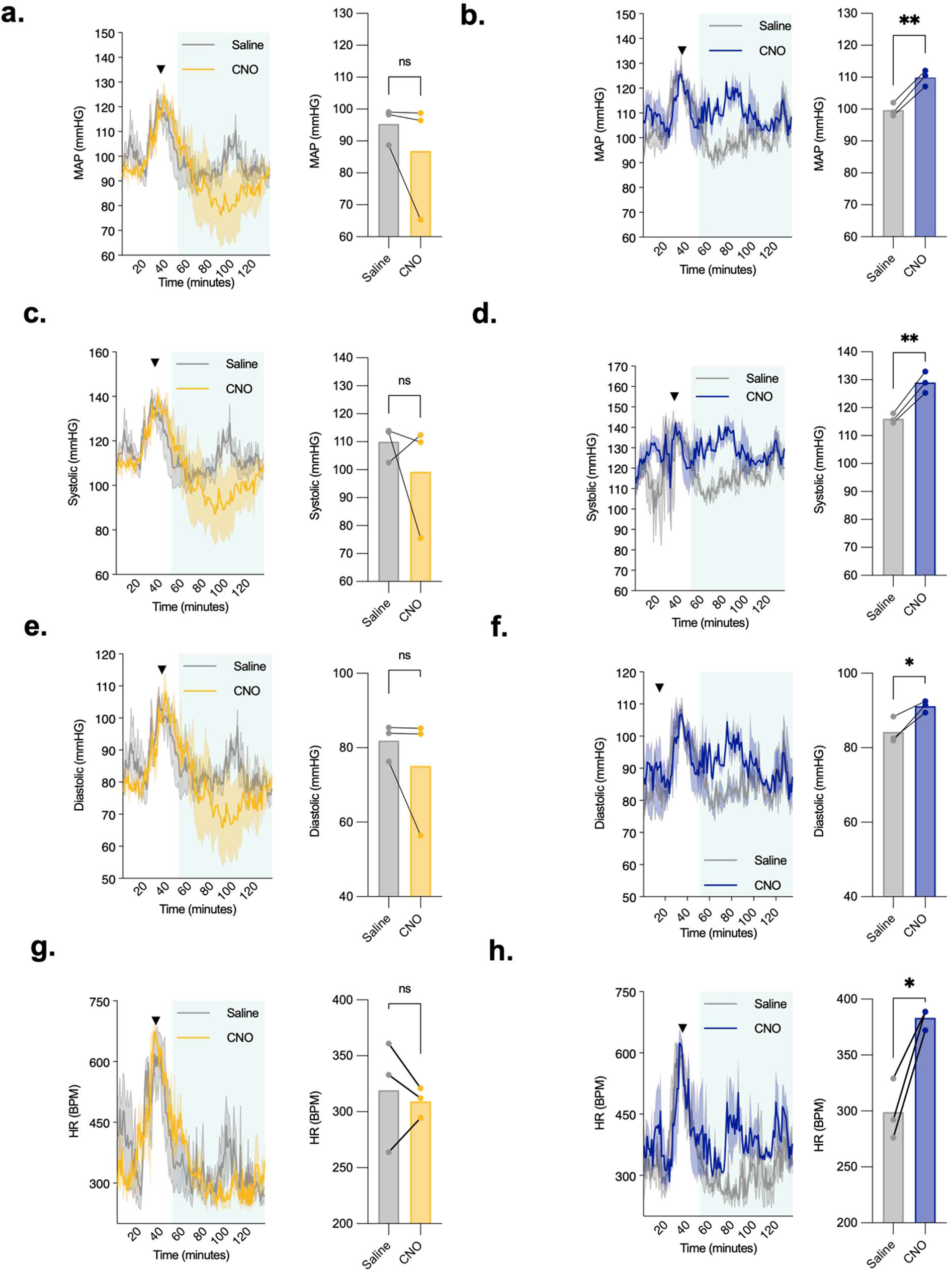
Modulation of cardiovascular function by ARC and DMH Lepr^Glp1r^ neurons. Shown are mean arterial pressure (MAP; a, b), systolic BP (c, d), diastolic BP (e, f), and HR (g, h) prior to and following the injection of saline (grey lines) or CNO in male Lepr^Glp1r-ARC-Dq^ (yellow; n=3) or Lepr^Glp1r-DMH-Dq^ mice (blue; n=3) mice. Left panels show continuous measurements; injection (t=40 minutes) is indicated by the arrowhead. Right panels show mean values (-/+ SEM) for 60-120 minutes (60 minutes, starting 20 minutes after the injection). *p< 0.05 and **p< 0.01 by Paired *t* test.

## Discussion

While Lepr^Glp1r^ neurons were previously shown to suppress food intake [19], our present findings reveal that Lepr^Glp1r^ neurons also mediate the control of cardiovascular function (HR and BP) by leptin. Furthermore, we discovered that the overall population of Lepr^Glp1r^ neurons is composed of two transcriptionally, anatomically, and functionally distinct subpopulations of cells. One of these subpopulations (ARC-residing Lepr^Glp1r-1^ neurons) suppresses food intake, while the other (DMH Lepr^Glp1r-2^ cells) controls cardiovascular function and other parameters associated with the control of SNS outflow. Hence, these results enable the dissociation of transcriptionally-defined *Lepr* neuron populations that control food intake from those that modulate cardiovascular function.

The neuron population referred to here as Lepr^Glp1r^ cells has also been studied as an ARC/DMH neuron population defined by the expression of *Trh* or *Bnc2*, or as a DMH population marked by *Glp1r* expression [14–16]. These GABAergic *Lepr* neurons directly innervate and inhibit *Agrp* neurons, suggesting that Lepr^Glp1r^ cells may mediate the suppression of food intake in part by restraining *Agrp* neuron activity [36]. Interestingly, our interrogation of leptin-regulated transcriptomic profiles for hypothalamic neurons, combined with GWAS data, suggested potential additional roles for Lepr^Glp1r^ neurons in the control of BP by leptin. Indeed, selective disruption of *Lepr* in the Lepr^Glp1r^ population lowered BP, even in the presence of pronounced hyperphagic obesity, which usually would be expected to increase BP [25].

Our molecular and spatial analyses identified two transcriptionally distinct subpopulations of Lepr^Glp1r^ neurons, each with unique gene expression profiles and anatomical distributions. To interrogate the circuitry of ARC vs DMH Lepr^Glp1r^ neurons in isolation (without simultaneously examining other intermingled *Lepr* neuron populations), we developed and employed several systems with which to modulate Lepr^Glp1r^ neurons specifically. We began by generating *Glp1r^Flp^* mice to enable the specific manipulation of Lepr^Glp1r^ neurons using intersectional recombinase techniques in Lepr^Cre^;Glp1r^Flp^ mice. We then employed our previously described tTARGIT system in Lepr^Cre^;Glp1r^Flp^ mice to express ChR2-YFP specifically in anatomically defined Lepr^Glp1r^ neurons, enabling us to examine downstream projections from ARC or DMH Lepr^Glp1r^ neurons. Additionally, we developed a dual-recombinase-dependent retrograde tracing system (the RADRR tracing system) to map the inputs to Lepr^Glp1r^ neurons specifically. The circuit mapping enabled by these tools defined unique neuronal inputs and outputs for each of the two Lepr^Glp1r^ subpopulations, suggesting their distinct functions.

Furthermore, gain of function experiments using dual-recombinase/tTARGIT-dependent chemogenetic activation enabled us to determine that ARC Lepr^Glp1r^ neurons suppress feeding without significantly altering energy expenditure, locomotor activity, BAT thermogenesis, and BP/HR – consistent with their direct projections to feeding-related hypothalamic regions. Similarly, we were able to show that activating DMH *Lepr^Glp1r^* neurons augments SNS-related parameters (energy expenditure, locomotor activity, BAT thermogenesis, or BP/HR) without altering food intake-consistent with their projections to the RPa, which plays important roles in controlling SNS outflow [7, 26]. These findings the reveal the likely molecular identity of the DMH *Lepr* cells (Lepr^Glp1r-DMH^ neurons) previously suggested to increase BP in response to leptin [8].

The hM3Dq-mediated activation of DMH Lepr^Glp1r^ neurons augments parameters affected by SNS outflow rapidly (within an hour), while the activation of ARC Lepr^Glp1r^ neurons detectably suppresses food intake only after longer times (4-6 hours after CNO treatment). The slower time course for the modulation of feeding than SNS outflow could reflect the difficulty of inhibiting the feeding effects of *Agrp* neurons when they are fully activated by orexigenic stimuli – potentially exacerbated by the unilateral nature of our Lepr^Glp1r^-focused manipulations with hM3Dq. While it is possible that bilateral activation of ARC Lepr^Glp1r^ neurons would inhibit *Agrp* neurons more uniformly, leading to a more rapid attenuation of feeding, the difficulty of confining injections to the ARC (or DMH) without overflow into the DMH (or ARC) rendered bilateral targeting impractical.

Because both subpopulations of Lepr^Glp1r^ neurons project to *Agrp* neurons, their distinct effects might stem from their modulation of different subsets of *Agrp* neurons. Indeed, *Agrp* neurons segment into multiple distinct populations of neurons in some single-cell analyses [27, 28], and individual *Agrp* neurons exhibit single projection targets [11, 29–31], consistent with the notion that it might be possible to control different aspects of physiology and behavior by targeting different groups of *Agrp* cells. Additionally, or alternatively, the differing effects of ARC and DMH Lepr^Glp1r^ neurons might result from their other (distinct) projection targets.

Therapeutically targeting subpopulations of Lepr^Glp1r^ neurons might enable the selective modulation of feeding or cardiovascular parameters, potentially improving the efficacy and safety of medicines to treat cardiometabolic illness. Further studies employing loss-of-function approaches and evaluating the function of these neurons under pathophysiological conditions will help pinpoint therapeutic opportunities and further clarify the role for each Lepr^Glp1r^ neuron subpopulation in homeostatic regulation. While GLP1RAs activate both Lepr^Glp1r^ neuron subpopulations, it is possible that other approaches could enable the selective activation (or deactivation) of a single Lepr^Glp1r^ subpopulation.

Interestingly, genes in hypertension-associated GWAS loci [17] enriched specifically with genes whose expression is regulated by leptin in Lepr^Glp1r^ neurons, rather than with the transcriptome of these neurons at baseline. This link, together with the finding that Lepr^Glp1r^ neurons mediate the control of BP by leptin and previous data linking loss-of-function mutations in *LEPR* and *MC4R* with reduced BP [[Zorn, 2025 #2][8], suggests a potential role for Lepr^Glp1r^ neurons in the hypertension that accompanies obesity/hyperleptinemia. Given that DMH (but not ARC) Lepr^Glp1r^ neurons modulate BP and other aspects of cardiovascular function, we predict that the leptin-regulated transcriptional response of Lepr^Glp1r-2^ neurons would correlate more strongly with hypertension GWAS genes than leptin-regulated genes in Lepr^Glp1r-1^ neuron genes. Unfortunately, leptin-regulated snRNA-seq data sufficiently powered to subset the leptin response of the two Lepr^Glp1r^ neuron subpopulations do not exist at present.

In summary, our findings reveal roles for Lepr^Glp1r^ neurons in the augmentation of cardiovascular function, in addition to the suppression of food intake. We also identify two distinct populations of Lepr^Glp1r^ neurons that mediate specific physiological and behavioral outputs. Hence, this study provides a more precise framework for understanding the hypothalamic control of metabolic and cardiovascular function by leptin and opens avenues for more targeted therapeutic interventions.

## Methods

### Animals

All procedures performed on animals were approved by the University of Michigan Institutional Committee on the Care and Use of Animals (approval number PRO00012645) and in accordance with AAALAC and NIH guidelines. All mice were provided with water *ad libitum* and housed in temperature-controlled rooms on a 12-hour light-dark cycle. All mice were provided *ad libitum* access to standard chow diet (LabDiet 5L0D) unless otherwise noted.

Mice subjected to MBH snRNA-seq are described in [18]. All other mice were bred in our colony in the Unit for Laboratory Animal Management at the University of Michigan. Mice were weaned at 21 days and group housed with same-sex littermates fed the appropriate diet. *Glp1r^cre/cre^;Lepr^Flox/Flox^* (Lepr^Glp1r^KO) and *Lepr^cre/cre^;Rosa26^Sun1-sfGfp/Sun1-sfGfp^* (Lepr^Sun1-sfGfp^) mice were as previously described.

We generated *Glp1r^Flpo^* mice using CRISPR/Cas9-directed gene targeting in the MDRC Molecular Genetics Core. We targeted the 3’-end of the coding sequences of the final exon of *Glp1r* using a pair of sgRNAs targeting the sequences ccacaggaggaccggcactg gctgggtggccatcccaggt. To replace the *Glp1r* STOP codon with sequences fusing 2A peptide plus a codon-optimized FlpO recombinase, the sgRNAs were co-injected into fertilized mouse embryos along with the megamer editing template 5’-ctgcagcatgaagcccctcaagtgtcccaccagcagcgtcagcagtggggccacagtgggcagcagcgtgtacgcagccacctgccagagttcctacagcGGTTCTGGCGCAACAAATTTCTCACTCTTGAAACAGGCAGGAGATGTCGAAGAGAACCCAGGACCTATGCCCAAAAAGAAGAGGAAGGTCTCCCAATTCGATATCCTTTGCAAAACTCCACCTAAGGTGTTGGTGAGACAATTCGTAGAGAGGTTTGAAAGGCCCAGTGGAGAAAAGATCGCTTCATGTGCCGCTGAACTGACCTACTTGTGTTGGATGATTACACACAATGGAACCGCTATAAAGAGGGCCACTTTCATGAGTTACAATACAATAATTTCTAACTCCTTGAGCTTCGACATAGTCAATAAGTCCCTCCAGTTCAAATACAAAACTCAGAAAGCTACCATACTCGAGGCAAGCCTCAAGAAACTGATACCCGCCTGGGAGTTCACTATCATCCCCTACAACGGTCAAAAACACCAGTCAGATATAACAGACATAGTATCCAGTCTGCAACTCCAGTTCGAGAGCAGCGAGGAAGCCGACAAAGGCAATAGTCACAGCAAGAAAATGCTTAAAGCATTGTTGTCCGAGGGGGAAAGTATTTGGGAAATCACTGAAAAGATCCTGAACAGTTTCGAATATACATCCCGCTTTACTAAAACTAAAACACTTTACCAGTTCCTCTTTCTTGCCACATTTATAAATTGCGGTCGATTCTCAGACATAAAGAATGTAGACCCCAAATCCTTCAAGCTCGTTCAAAATAAATATTTGGGGGTCATTATTCAGTGCCTGGTGACAGAGACCAAGACCTCAGTTTCACGGCATATTTATTTCTTCTCAGCTAGAGGGCGGATAGATCCTCTTGTATATCTCGACGAATTCCTTCGGAATAGTGAACCAGTCTTGAAGCGAGTTAATCGCACCGGCAACTCAAGTAGTAACAAACAAGAATATCAGCTCTTGAAGGACAATCTTGTTCGATCTTATAACAAAGCACTGAAGAAGAATGCACCCTATCCTATCTTTGCAATAAAAAATGGGCCTAAAAGCCATATCGGACGGCACCTCATGACTAGCTTTCTTAGCATGAAAGGGCTGACAGAGCTGACTAATGTCGTAGGCAACTGGAGCGATAAAAGAGCCAGTGCCGTTGCACGGACAACTTACACCCATCAGATTACTGCTATACCAGACCACTATTTCGCCCTGGTTTCTAGGTACTACGCTTATGACCCAATATCAAAAGAGATGATTGCACTGAAGGATGAGACTAACCCCATAGAAGAATGGCAACATATTGAGCAACTGAAGGGAAGCGCCGAGGGATCCATTCGATATCCCGCTTGGAACGGTATCATTTCACAGGAAGTTCTCGACTACTTGAGCTCTTACATTAACAGGCGCATCTAATGAcctcctgtggtccttgcttctggctgggtggccatcccaggtCATagagatcctggggatagggaatgtgaaggacacaggcacaccacacacac-3’.

Embryo injections were performed by the University of Michigan Transgenic Animal Core. Fertilized embryos were implanted into pseudopregnant dams. Resultant pups were screened for the correctly-targeted insertion of 2A-Cre sequences by PCR. We subsequently amplified and sequenced genomic sequences spanning from outside of the 5’ end to outside of the 3’ end of the editing template. Positive animals were bred to C57Bl6/J mice and the resultant pups were rescreened and resequenced prior to propagation.

*Glp1r^Flpo^* mice were bred to *Lepr^cre^* mice from our colony to generate *Lepr^Cre/Cre^;Glp1r^Flpo/Flpo^* (*Lepr^Cre^;Glp1r^Flp^*) mice and to *Rosa26^LSLGfp-L10a^* [32]and *Rosa26^FSFtdTomato^* (*Gt(ROSA)26Sor^tm65.2(CAG-tdTomato)Hze^*/J; Jax Strain #032864) to generate Lepr*^Cre/Cre^_;_*Glp1r*^Flpo/Flpo^_;_*Rosa26*^LSLGfp-L10a/FSFtdTomato^* (Lepr^Gfp^;Glp1r^tdT^) reporter mice.

### Viral vectors

All AAVs were packaged into AAV8 capsids. tTARGIT AAVs and AAV-hSyn-DIO-hM3Dq have been described (Sabatini). To generate RADRR plasmids, rabies glycoprotein and TVAmcherry were individually PCR amplified using TRE-DIO-TVAmCherry-rG (Addgene plasmid # 166602) as a template. The resulting rabies glycoprotein amplicon was inserted into an Flp-dependent AAV plasmid backbone (AAV-hSYN1-fDIO-GFP) and the TVAmcherry amplicon was inserted into a Cre-dependent AAV backbone (AAV-hSYN1-DIO-hM3DqmCherry; Addgene plasmid #44361) using standard restriction enzyme and ligation approaches. Correct incorporation of the transgenes into the AAV backbone was confirmed by Sanger sequencing. All AAVs were packaged and EnvA-psuedotyped Rabies^ΔG-Gfp^ was produced by the Michigan Diabetes Research Center/University of Michigan Viral Vector Core.

### Analysis of KO mice

Body weights were taken from *Lepr^Flox/Flox^* and *Glp1r^Cre^;Lepr^Flox/Flox^* at 9 weeks of age. BP and HR measures were recorded for ten continuous seconds every ten minutes. Average BP and HR were taken during the first 30 minutes of the onset of the light cycle in female mice collected from DSI BP Analysis software. Significance for bodyweight and cardiovascular measures were determined via Welche’s *t* test.

### Leptin injection data and GWAS enrichment analysis

Leptin injection data from [18] was used for the following analyses; please refer to that paper for information on mice and single-cell data generation. To assess exogeneous leptin-driven shifts in cellular composition between ip leptin and saline treatment mice, the Seurat object was converted to a Milo [33] object and neighborhood differential abundance (DA) testing was performed using TMM normalization and graph-overlap FDR weighting. A design matrix (∼0 + group) was used to test both genotype-by-treatment interaction and within-genotype treatment contrasts. Neighborhoods were annotated by dominant predicted cell type and experimental group. Neighborhood-level pseudobulk expression profiles were generated by summing raw RNA counts across cells within each Milo neighborhood. Gene counts were analyzed using DESeq2 [34] with a design of ∼ predicted.celltype + group. Lowly expressed genes were filtered (≥5 counts in ≥20 neighborhoods), and variance-stabilizing transformation was applied. To integrate systolic BP GWAS signal [17] mouse genes were mapped to human orthologs and converted to Entrez identifiers using org.Hs.eg.db, retaining unique mappings.

Average neighborhood expression values were computed per gene and used as covariates in MAGMA [35] gene-set analysis. SNP-wise gene statistics were first generated using MAGMA with a European 1000 Genomes reference panel [36]. Gene-set analysis was then performed using neighborhood-level gene expression covariates and a one-sided enrichment model. Resulting neighborhood GWAS enrichment statistics were multiple-testing corrected [37] and integrated with Milo-derived neighborhood annotations and group composition for downstream visualization and regression analyses.

### Single-Nucleus RNA Sequencing

We euthanized Lepr^Sun1-sfGfp^ mice (n=4, mixed sex) with isoflurane, then decapitated them. We dissected the brain and sectioned it into 1 mm thick coronal slices at the mediobasal hypothalamus using a brain matrix, then flash-froze samples in liquid nitrogen.

We homogenized frozen tissue in Lysis Buffer (EZ Prep Nuclei Kit, Sigma-Aldrich) supplemented with Protector RNAase Inhibitor (Sigma-Aldrich). We filtered the homogenate through a 30 μm MACS strainer (Miltenyi) and centrifuged it at 500 rcf for 5 minutes at 4°C to pellet the nuclei. Next, we resuspended the nuclei in wash buffer (10 mM Tris, pH 8.0; 5 mM KCl; 12.5 mM MgCl2; 1% BSA; RNAse inhibitor), then repeated filtration and centrifugation. We stained intact nuclei with propidium iodide (Sigma-Aldrich), then sorted them using a MoFlo Astrios Cell Sorter to collect GFP+ and PI+ nuclei for LepRb-Sun1 experiments. After sorting, we centrifuged nuclei at 100 rcf for 5 minutes at 4°C and resuspended them in wash buffer to achieve a concentration of 750–1,200 nuclei/μL. We added RT mix to capture approximately 10,000 nuclei, then loaded the mixture onto a 10× Chromium Controller chip. We prepared libraries using the Chromium Single Cell 3′ Library and Gel Bead Kit v3, Chromium Chip B Single Cell kit, and Chromium i7 Multiplex Kit as per manufacturer’s instructions. Libraries were sequenced on an Illumina NovaSeq 6000 (paired-end, 150 nt read lengths).

### snRNA-seq Data Analysis

We mapped FASTQ files to the reference genome (Ensembl GRCm38) using CellRanger 6.12 to generate count matrices, then analyzed data using Seurat in R. We retained nuclei with at least 800 detected genes and less than 6% mitochondrial gene content. We used DoubletFinder to identify and remove suspected doublets. After excluding doublets, we identified variable features per sample with FindVariableFeatures, then scaled gene expression. We used PCA on variable features to assess batch effects, then performed UMAP projection and clustering using the FindNeighbors and FindClusters functions. We integrated data across samples for joint analysis.

To assign cell types, we projected cell type labels from a published snRNA-seq dataset (GSE87544) using Seurat PCA. We selected neuronal nuclear clusters and excluded thalamic nuclei using markers identified in previous thalamic sequencing datasets.

Hypothalamic nuclei then underwent a final round of quality control based on mitochondrial gene content, excluding nuclei with less than 0.15% mitochondrial content. We clustered the remaining nuclei at a resolution of 0.55 and named clusters based on unbiased gene markers identified via FindClusterMarkers.

### Differential Gene Expression Between Lepr^Glp1r-1^ and Lepr^Glp1r-2^ Clusters

We used Seurat’s FindMarkers to compare the two clusters, setting the log fold change threshold at 2. For visualization, we included genes with p-values < 0.001 and expression in at least 20% of nuclei.

### Spatial transcriptomics visualization

The spatial transcriptomics analyses were based on re-analyses of data generated for [18]. For the present study, single-section Xenium objects were loaded individually and cell type classification was performed on nucleus-level transcript counts. *Lepr^Glp1r^* were classified as (Lepr+/Glp1r+/Trh+/Ebf1−) (Lepr^Glp1r-1^/ARC), (Lepr+/Glp1r+/Ebf1+/Trh−) (Lepr^Glp1r-2^/DMH), or quadruple-positive (Lepr+/Glp1r+/Trh+/Ebf1+) (Lepr^Glp1r-2^/DMH), requiring a minimum of one transcript count per gene per cell for a gene to be considered expressed. All remaining cells were classified as background. Spatial visualization was performed using the ImageDimPlot function from the Seurat v5 package in R [Hao, 2021 #199;Dylan M Belmont-Rausch 1, 2026 #78], rendering nucleus segmentation boundaries colored by subpopulation identity. Individual transcript molecules for the landmark genes *Agrp*, *Sim1*, and *Fezf1* were overlaid as single-molecule dot plots (5,000 molecules sampled per gene, alpha=0.1) to provide anatomical reference for the arcuate nucleus (ARC; *Agrp*), dorsomedial hypothalamus (DMH; *Sim1*), and ventromedial hypothalamus (VMH; *Fezf1*). Anatomical region boundaries were manually annotated based on landmark gene distributions and anatomical reference sections at Bregma -2mm. Figures were exported as vector graphics using the Cairo PDF device at 300 DPI.

### FOS Studies

We performed FOS activation studies in singly housed Lepr^CreL10GFP^;Glp1r^FlpTDT^ and Lepr^CreSun1-GFP^;Glp1r^FlpTDT^ reporter mice. All animals were fasted for 16 hours beginning at the onset of the dark cycle. We injected vehicle, leptin (5 mg/kg, IP) or semaglutide (10 nmol/kg, IP) 2 hours prior to perfusion and tissue collection.

Following treatment, we sedated the mice with isoflurane and perfused them with formalin to fix the brain tissue. After decapitation, we post-fixed brains in formalin for 4 hours, then transferred them to 30% sucrose. We sectioned hypothalamic tissue into 30 μm slices at −40°C. We stained sections for GFP (*Lepr* neurons) and FOS. *Glp1r* neurons were detected via endogenous tdTomato fluorescence. We imaged FITC, mCherry, and Cy5 channels at 20x magnification.

To quantify FOS colocalization with Lepr^Glp1r^ neurons we analyzed images from each channel and region using CellProfiler to identify and quantify FOS (20–60 pixels/object), GFP (15–60 pixels/object), and tdTomato (15–70 objects/section) positive cells then used the data frames containing the pixel coordinates for objects in each channel to determine where the objects overlapped. We quantified the number of Lepr^Glp1r^ cells per section by counting GFP and TDT objects with centers within 25 pixels distance to one another. To assess FOS overlap with Lepr^Glp1r^ cells, we identified FOS, GFP, and TDT objects whose centers were within 25 pixels of one another.

### Stereotaxic Viral Injections

We anesthetized animals with isoflurane (induction: 5%, maintenance: 2.5%), placed them on a heating pad, and secured them in a stereotaxic frame (David Kopf Instruments). We administered carprofen (5 mg/kg) and sterile saline subcutaneously. After exposing the skull and identifying bregma, we targeted specific hypothalamic regions with a glass pipette: caudal ARC (Y: −1.4, X: −0.3, Z: −5.82), vDMH (Y: −1.5, X: −0.3, Z: −5.2), and TU (Y: −1.6, X: −0.9, Z: −5.5). We drilled at target coordinates and delivered virus (volume specified below) at the appropriate Z depth over 2 minutes, leaving the pipette in place for 3 minutes post-injection. We slowly withdrew the pipette over 3–5 minutes, sutured the incision, and monitored animals daily for 7 days, administering carprofen (5 mg/kg) IP 24 hours after surgery.

*Rabies Retrograde Tracing*. We performed two intracranial surgeries in Lepr^Cre^;Glp1r^Flp^ mice (8 weeks–10 months old). First, we co-injected the two RADRR helper viruses (AAV8-hSYN1-DIO-TVA-mCherry and AAV8-hSYN1-fDIO-rG) into the vDMH (100 nL) or rostral ARC (50 nL). After at least three weeks later, we injected 50 nL of pseudotyped Rabies^ΔG-GFP^ [22] into the same site. Five days post-injection, we processed brains for immunohistochemistry. Controls included *Lepr^Cre^* or *Glp1r^Flp^* mice receiving identical treatments.

*Anterograde Tracing.* We used the tTARGIT system, which utilizes two viruses to mediate the Cre-plus Flp-dependent expression of ChR2-YFP in Lepr^Glp1r^ neurons. One virus (AAV-hSYN1-fDIO-tTA) drives tetracycline transactivator expression, and the second (rAAV8-TRE-DIO-ChR2-EYFP) drives Cre-dependent ChR2-YFP expression in the presence of tTA. We co-injected the two viruses unilaterally into *Lepr^Cre^;Glp1r^Flp^* mice; some *Lepr^Cre^* only or *Glp1r^Flp^* only mice were also injected as controls. A minimum of three weeks following surgery, we perfused the mice and processed their brains for immunohistochemical analysis.

*DREADD (hM3Dq) expression.* We used the tTARGIT system to express hM3Dq in Lepr^Glp1r^ neurons in the DMH and ARC of Lepr^Cre^;Glp1r^Flp^ mice (8–10 weeks old) by unilaterally co-injecting AAV-hSYN1-fDIO-tTA and AAV1-TRE-DIO-hM3Dq-mCherry. Animals were allowed to recover for at least three weeks following surgery before study. At study completion, we injected CNO (1 mg/kg, IP) 90 minutes before euthanasia and perfusion, we then sectioned post-fixed brains into 30 μm slices for post hoc analysis.

Animals displaying off-target or insufficient hM3Dq-mCherry expression were excluded from analysis.

### Functional Studies in hM3Dq-expressing Animals

*Metabolic Cage Studies.* Following recovery from surgery, we housed DMH-targeted and ARC-targeted animals singly for one week prior to indirect calorimetry in a Promethion System (Sable Systems International) to measure energy expenditure, activity, and food intake over 8 days. Days 0–2 served as baseline (twice-daily saline IP injections, 9 am and 4 pm); days 3–5 included twice-daily CNO injections; and days 6–7 were washout days. We measured VO_2_, VCO_2_, and locomotor activity using the system’s open-circuit calorimetry and optical beam monitoring. Animals were weighed and placed individually in mouse cages (Model 3721; 21 × 37 × 14 cm) with *ad libitum* access to food and water. We calculated energy expenditure from VO_2_ and VCO_2_.

For fasting/refeeding crossover studies, DMH-targeted or ARC-targeted mice were housed in a CLAMS system (Columbus Instruments). After baseline recordings (days 0–1), all mice received CNO or saline at 3:40 pm prior to, and at the end of, overnight fasting at 5:00 pm and 9:00 am on days 1, 2, 4, and 5. We weighed and injected animals according to body weight × 10. Days 6–7 were washout days.

*Temperature Monitoring*. We surgically implanted temperature Transponders (IPTT-300, BMDS) subcutaneously in the intrascapular space (between BAT lobes, in direct contact with BAT) and later in the abdominal cavity (secured in the hepatic portal region) of hM3Dq-expressing animals. Incisions were closed with one suture, and animals recovered for 5 days before undergoing saline/CNO injections and temperature scans.

In week 1, we recorded temperature after IP saline or CNO injections; in week 2, injection types were crossed over. Temperature was recorded manually using DAS-7007S scanner (BMDS) without direct animal contact. Baseline temperature was measured at 10:00 am, followed by IP injection of 1 mg/kg CNO (0.1 mg/mL, volume based on body weight × 10). We measured temperature at 15, 30, 60, 120, 180-, 240-, 300-, and 360-minutes post-injection.

### Telemetric Monitoring of Cardiovascular Function

Lepr^Glp1r^KO and their controls, or hM3Dq-expressing mice underwent implantation of a cardiac telemeter device to directly measure cardiac function. Mice were anesthetized and then placed on a heating pad and monitored body temperature throughout surgery. Once adequate anesthesia was confirmed, we made a midline abdominal incision and separated the abdominal muscles. We implanted the body of the telemeter (DSI, Cat# HD-X02) in the peritoneal cavity. We tunneled the electrode leads subcutaneously to place the positive lead on the apex of the heart and the negative lead in the right upper thorax, securing them with a small suture. We closed both the muscle and skin layers with absorbable sutures. Mice recovered from anesthesia in a warmed chamber and received analgesia postoperatively. We monitored all animals daily for signs of pain or distress.

All animals were allowed at least 7 days to recover from telemeter implantation surgery. Lepr^Glp1r^KO animals and their controls were monitored continuously for 3 days. hM3Dq-expressing mice were acclimated to IP saline injections over 3 consecutive days. Following handling acclimation, we removed food at 8 pm and fasted the animals overnight until 10 am. At 10 am, we administered IP injections of either CNO or saline. We collected minute-by-minute measures of mean arterial pressure (MAP), systolic and diastolic BP, HR, and pulse pressure. Food was returned 2 hours after injection. After each session, animals rested for 5 days before undergoing the crossover procedure. To evaluate the change in cardiac function between DMH and ARC neuron activation, we calculated the minute-by-minute delta change for each parameter (saline vs. CNO) and plotted these traces for 15 minutes prior to injection and for 75 minutes post-injection.

To assess within-group differences, we calculated the AUC for the last 40 minutes of the first hour after injection. We used paired t-tests to compare saline and CNO conditions.

### Quantification and statistical analysis

All plotting and statistical analysis was performed using R 4.5.2 and Graph Pad Prism 10. Specific statistical tests are listed in the figure legends. Sample sizes and sex distribution are listed in Supplemental Table 1.

### Study approval

All procedures performed were approved by the University of Michigan Committee on the Use and Care of Animals and in accordance with Association for the Assessment and Approval of Laboratory Animal Care and National Institutes of Health guidelines.

## Author Contributions

AMD, DBR, AJT, DPO, PVS, MGM and THP designed the experiments and analyzed data. AMD, DBR, AJT. AC. FS, EH, BC, JMB, SH, ZW, and NQ acquired and analyzed data. AMD wrote the first draft of the manuscript in collaboration with MGM and THP. All authors reviewed and edited the manuscript. MGM and THP provided project oversight and acquired funding. MGM is the guarantor of the manuscript.

## Supplemental Figures and Legends

**Supplemental Figure 1.**
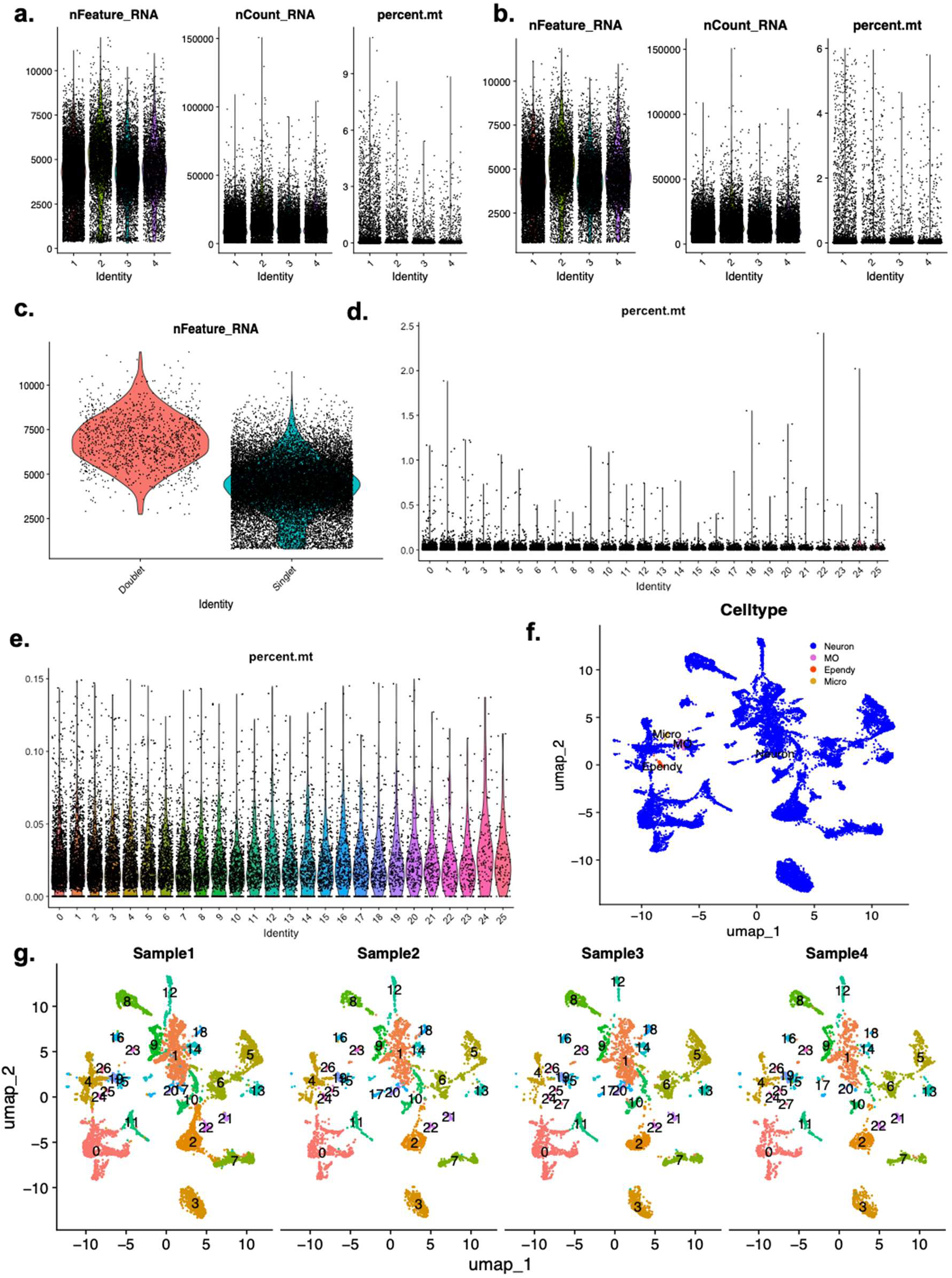
Initial QC of LepRb^Sun1-GFP^ mice sequencing dataset. (a) Distribution of raw features, feature counts and percent mitochondrial (mt) RNA across the four hypothalamic samples. (b) Distribution of features, feature counts and percent mt-RNA following the removal of all nuclei with fewer than 800 features and mtRNA greater than 6%. Feature number (c) for singlets and doublets. (d) Distribution of mtRNA between 0 and 2.5% across clusters defined by UMAP clustering of nuclei identified at singlets. (e) Distribution of mtRNA between 0 and 0.15%. (f) UMAP projection of nuclei colored by cell type and (g) UMAP projections of nuclei per sample following initial round of QC.

**Supplemental Figure 2.**
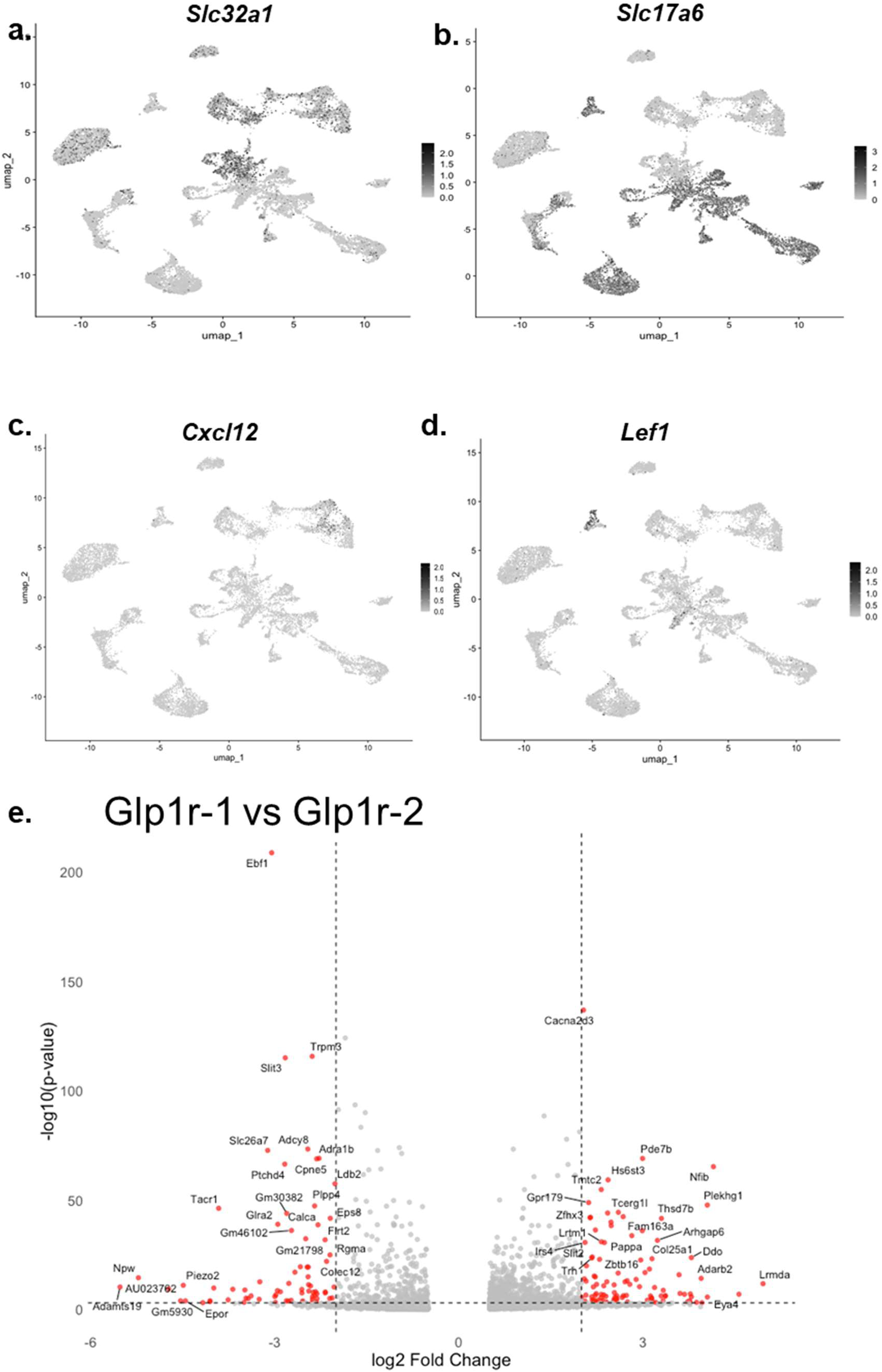
Distribution of marker genes across all populations of *Lepr* neurons. (a-d) and differential gene expression between Lepr^Glp1r-1^ and Lepr^Glp1r-2^ neuron populations. (a-d) Feature plots for *Slc32a1 (vGAT)* and *Slc17a6 (vGLUT2), Cxcl12, and Lef11* expression across neuronal nuclei post QC. (e) All genes differentially expressed between Lepr^Glp1r-1^ and Lepr^Glp1r-2^ clusters; p <0 .001 and Log2FC>2 for genes indicated in red.

**Supplemental Figure 3.**
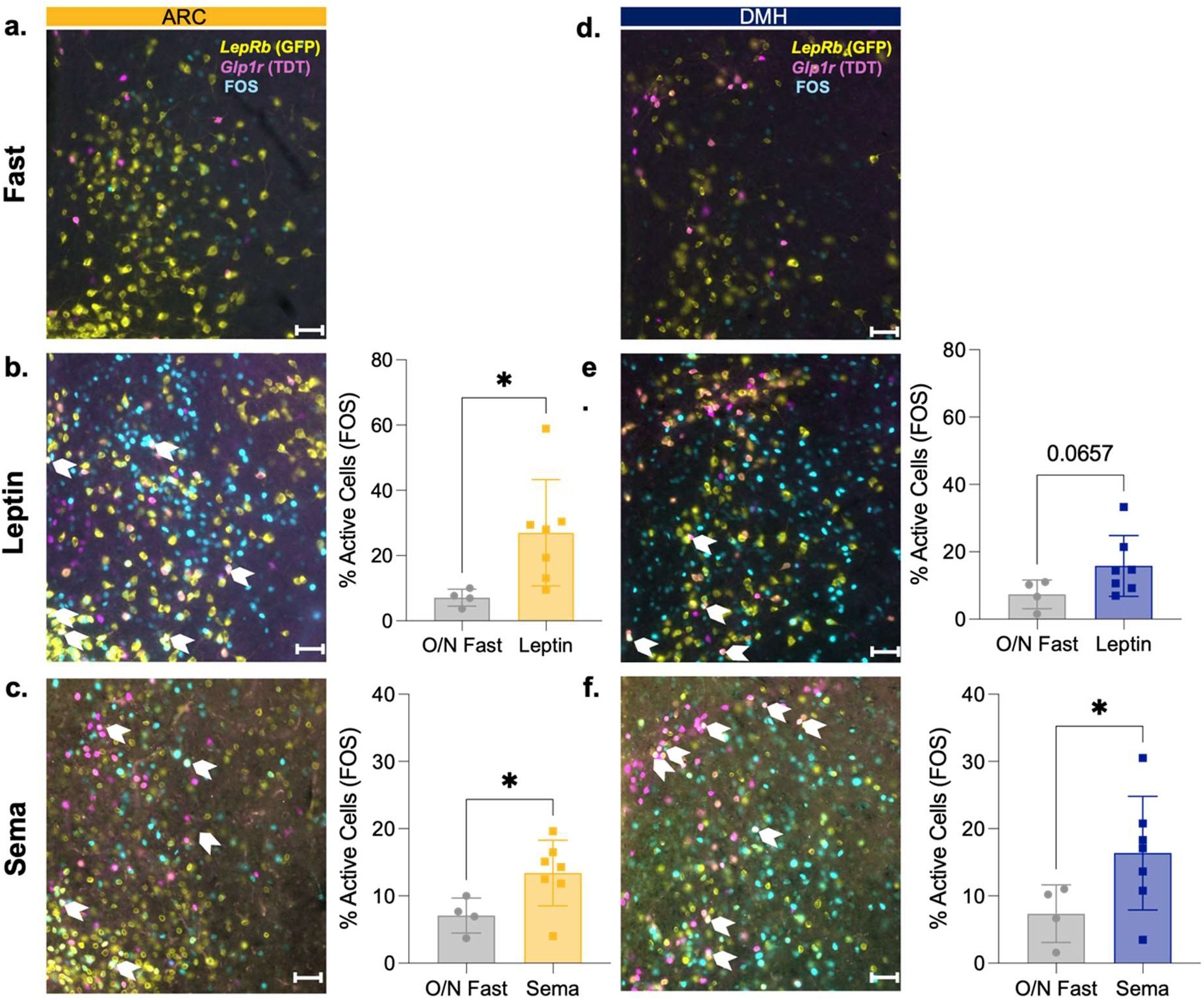
FOS accumulation in response to leptin and semaglutide in ARC and DMH Lepr^Glp1r^ neurons. Shown are representative images of the ARC (a-c) and DMH (e-g) of Lepr^GFP^;Glp1r^Tdt^ mice showing *Lepr* (GFP; yellow) and *Glp1r* (tdTomato; pink) and FOS-immunoreactivity (blue) cells. Images show each area following a 14 hour fast alone (a, d, n=4 males), followed by leptin treatment (1 mg/kg, IP)(b, e; n= 2 females and 5 males) or semaglutide (Sema; 10 nmol/kg, IP)(c, f; n= 3 females and 4 males). White arrows indicate cells positive for all three signals. Scale bars= 50 um. Left panels show quantification of FOS colocalized with GFP+tdTomato expressed as a percentage of total GFP+tdTomato cells. Mean +/-SEM are shown; *p< 0.05 by Welche’s *t* test.

**Supplemental Figure 4.**
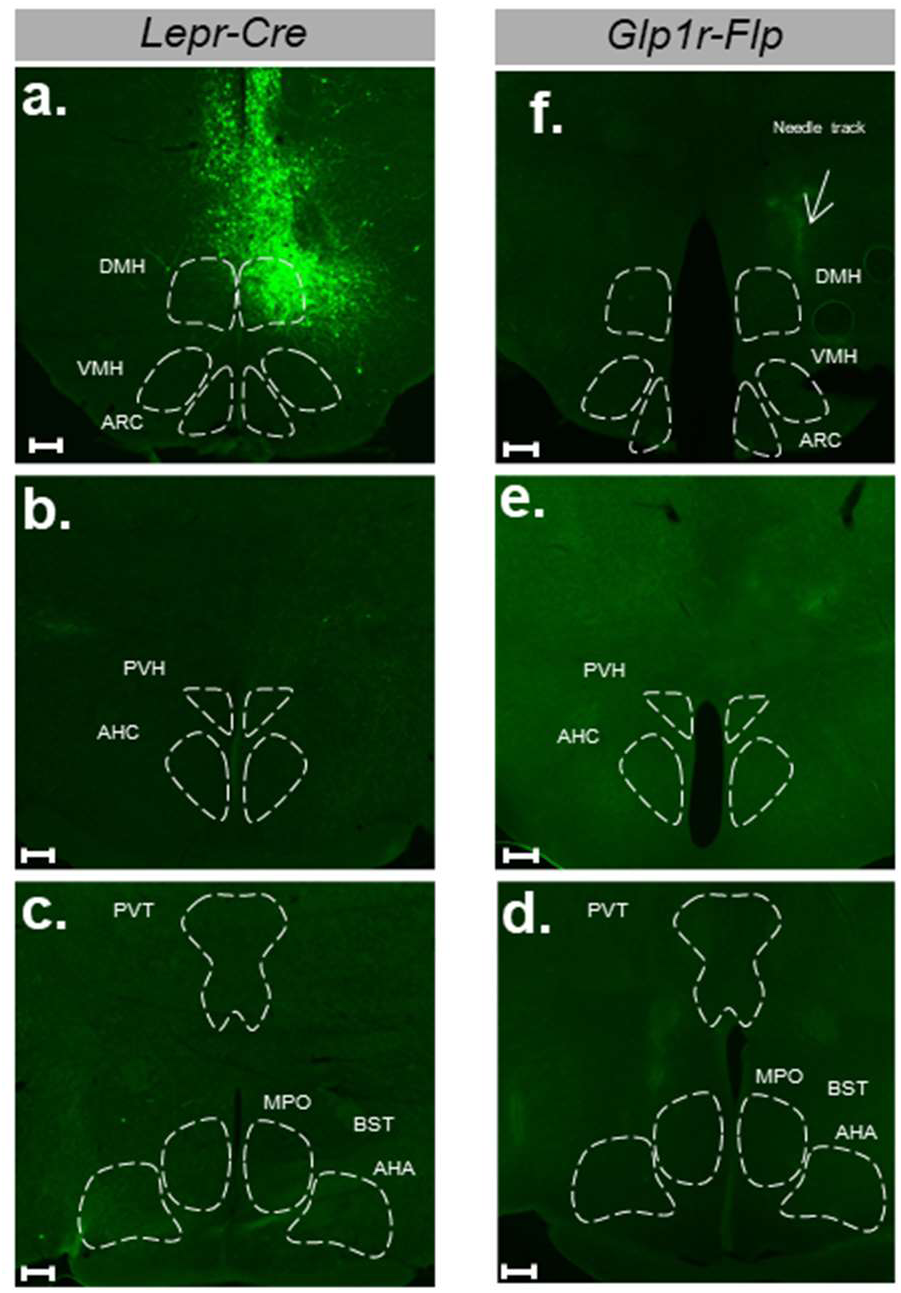
Lack of retrograde tracing following injection of RADRR tracing system into *Lepr^Cre^* only or *Glp1r^Flp^* only animals. Shown are representative images showing GFP (green) cell bodies following the intra-DMH injection of the RADRR tracing system in *Lepr^Cre^* mice (a-c; representative of n=2 males and 2 females) and *Glp1r^Flp^* (d-f; representative of n=2 females). Needle track for the DMH of *Glp1r^Flp^* mouse in (d) is indicated by a white arrowhead. Shown are the DMH injection site (a, d) and regions rostral to the DMH (b-c, e-f). Scale bar= 200 um.

**Supplemental Figure 5.**
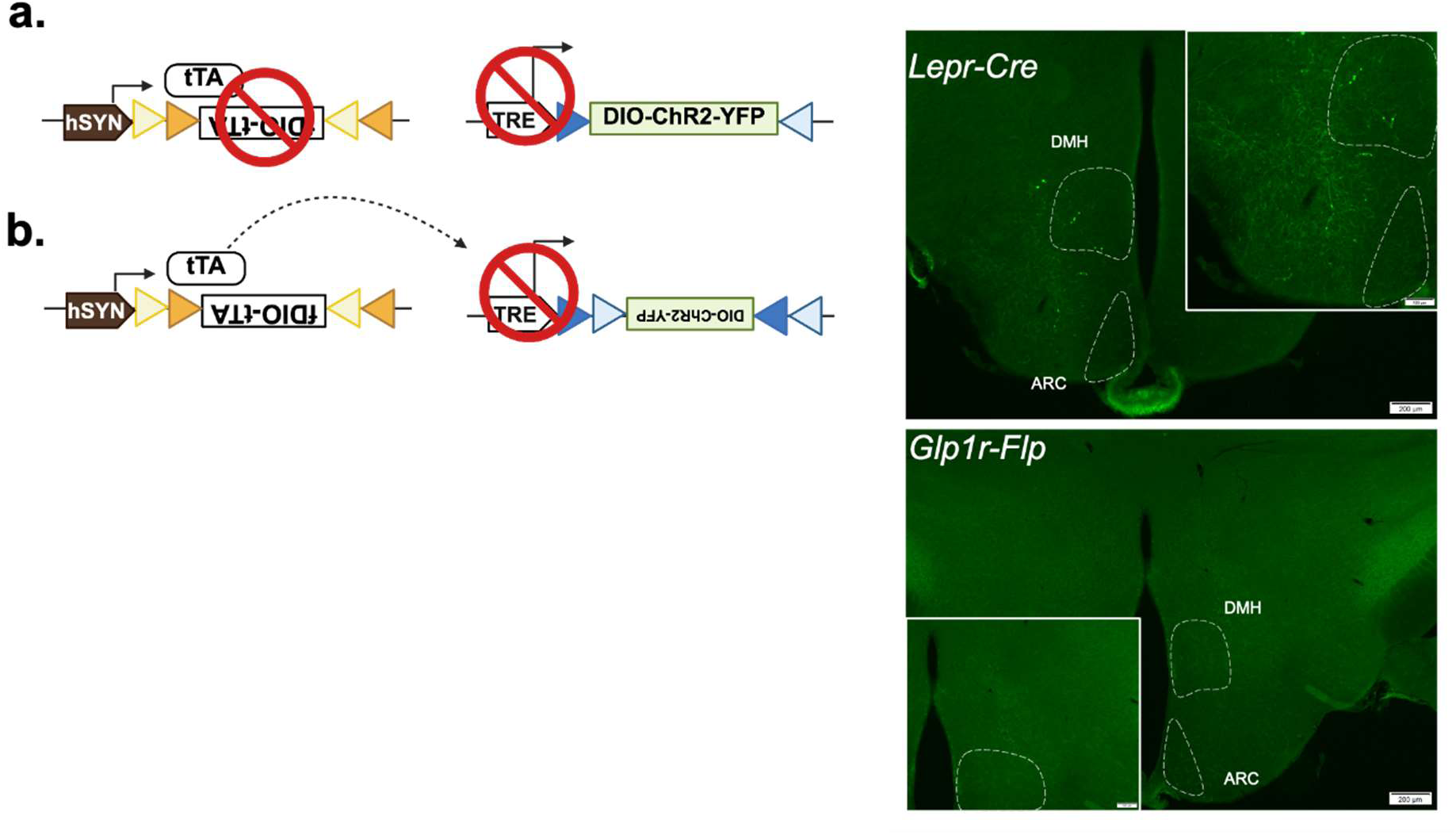
Lack of detectable ChR2-YFP following injection of the tTARGIT system into *Lepr^Cre^-* and *Glp1r^Flp^*-only controls. Shown are (left panels) schematics of the result of injecting the indicated tTARGIT system plasmids into recombinase expressing cells in *Lepr^Cre^* (a; representative of n=2 males and n=2 females) and *Glp1r^Flp^* (b; representative of n=4 females) mice, along with a representative image of the ChR2-eYFP (GFP; green) detection in targeted area (DMH, right panels) in each control. Scale bar in main images= 200 um. Scale bar in digital zoom inset= image 100um.

**Supplemental Figure 6.**
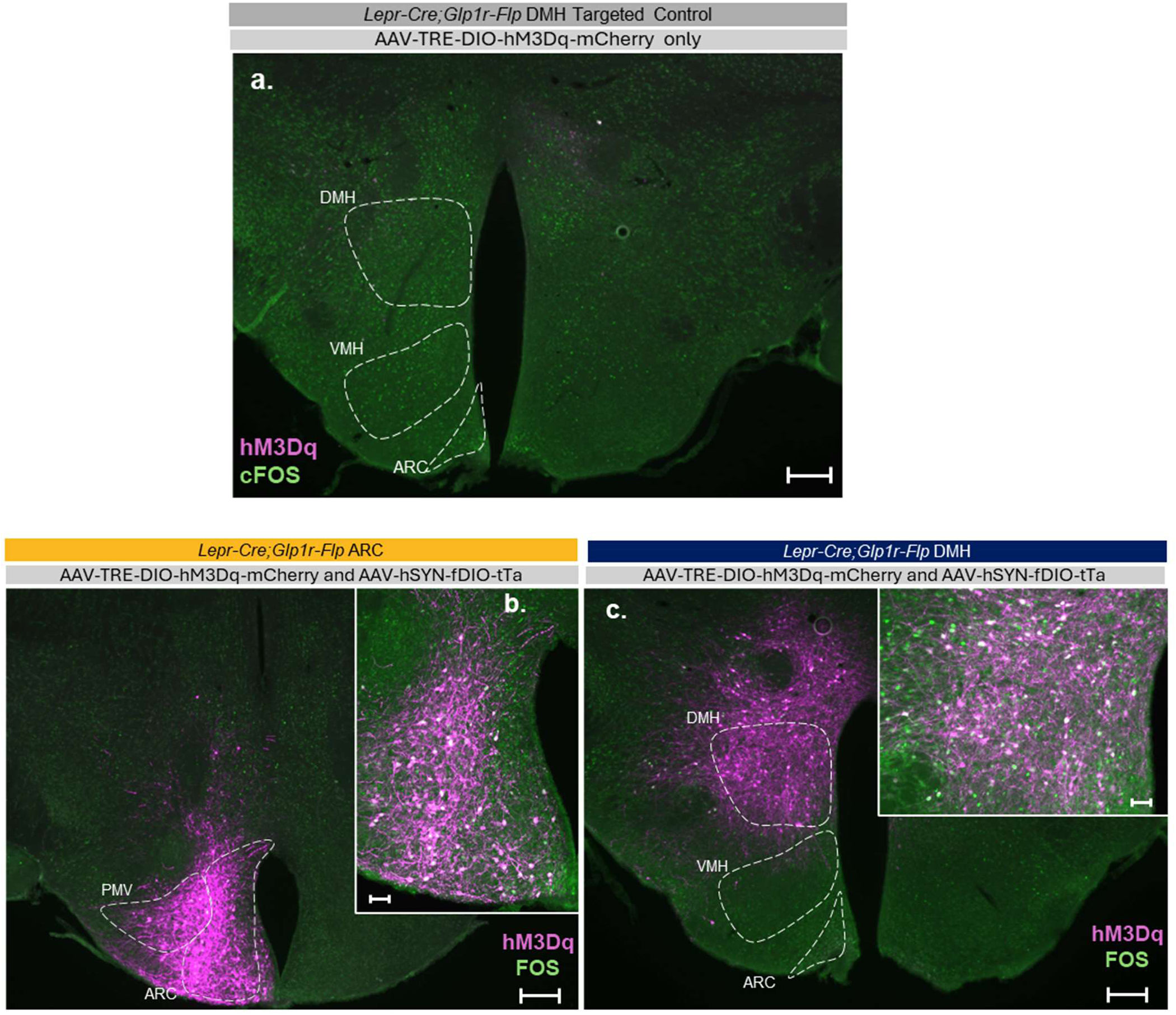
Controls for tTARGIT hM3Dq expression and activation of ARC and DMH Lepr^Glp1r^ neurons in correctly targeted injections. Shown are representative images of mCherry (hM3Dq; magenta) and FOS-immunoreactivity (green) in the hypothalami of *Lepr^Cre^*;*Glp1r^Flp^* mice injected into the DMH with AAV-TRE-DIO-hM3Dq-mCherry alone (top panel; representative of n=5 females and 3 males) or into the ARC (lower left; n=6 males, 5 females) or DMH (lower right, n=7 males, 4 females) with both AAV-TRE-DIO-hM3Dq-mCherry and AAV-hSYN-fDIO-tTa. Main panel scale bars= 200um, scale bars in digital zoom insets= 100um.

**Supplemental Figure 7.**
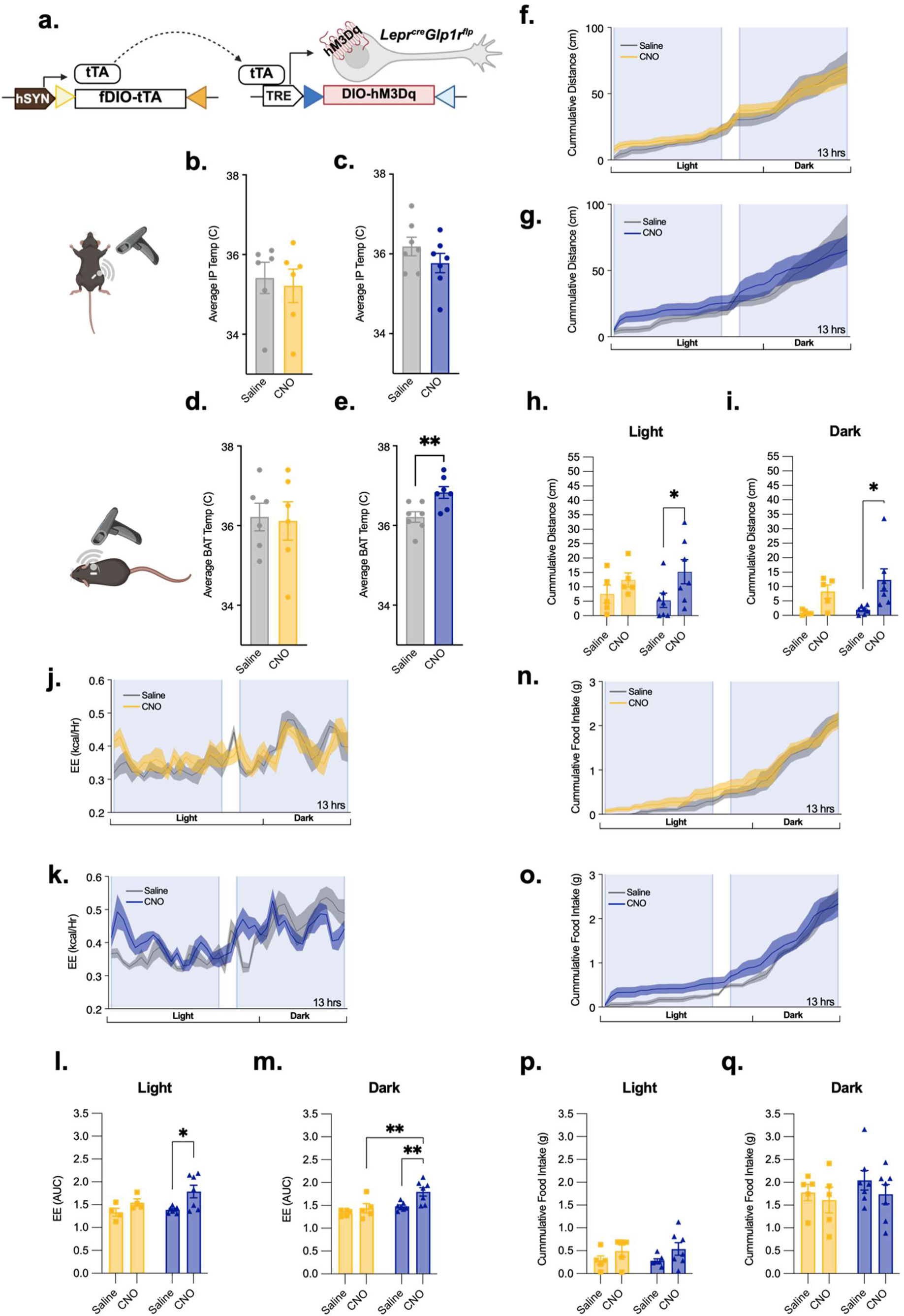
Modulation of food intake and energy expenditure by ARC and DMH Lepr^Glp1r^ neurons in male mice. (a) Schematic of the tTARGIT viral system used to express hM3Dq. (b, c) Average change in IP (core body) temperature (b, c) or intrascapular (BAT) temperature (d, e) over the 360 minutes following the activation of ARC (b, d; n=5) or DMH (c, e; n=7) Lepr^Glp1r^ neurons. Shown are averages +/-SEM; \**P* < 0.05 by Paired *t* test. (f-p) Continuous measurement of energy expenditure (f-i), cumulative locomotor activity (j-m), and cumulative food intake (n-q) following the injection of saline or CNO in Lepr^Glp1r-ARC-Dq^ (yellow; n=6) or Lepr^Glp1r-DMH-Dq^ mice (blue; n=7). (f, g, j, k, n, o) show continuous or cumulative measures of the indicated parameters over 13 hours; injections were performed at the beginning of the shaded areas. Graphs show summed values over the first hour (energy expenditure, locomotor activity) or six hours (food intake) following injection early in the light cycle (0900; h, l, p) or prior to the onset of the dark cycle (1600; i, m,q). Measurements were taken automatically using the SABLE metabolic cage system with 12 hour light and dark cycle. Comparisons were made between the third day of saline injections and first day of CNO injections. Bar graphs show averages +/-SEM; *p < 0.05, **p< 0.01 for the indicated comparisons by ANOVA with Fisher’s LSD post hoc test.

**Supplemental Figure 8.**
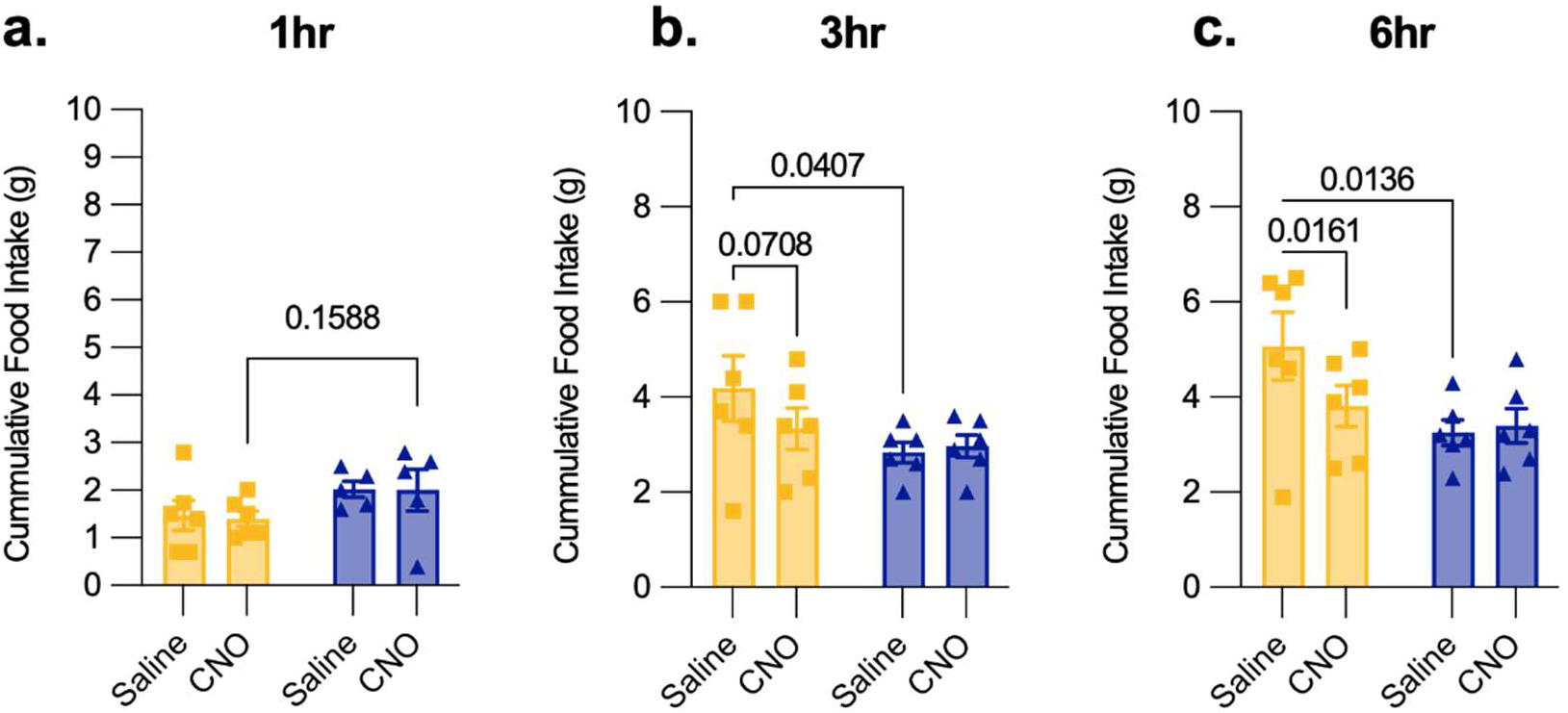
Effects of activating ARC and DMH Lepr^Glp1r^ neurons on food intake during refeeding following an overnight fast. Cumulative food intake following refeeding and IP injection (b: ARC; n=3 male and 3 female, c: DMH; n=3 male and 3 female) at 1 hour (a), 3 hours (b) and 4 hours (c). Bar graphs show averages +/-SEM; indicated comparisons by ANOVA with Fisher’s LSD post hoc test.

